# Preclinical efficacy and safety analysis of gamma-irradiated inactivated SARS-CoV-2 vaccine candidates

**DOI:** 10.1101/2020.09.04.277426

**Authors:** Gozde Sir Karakus, Cihan Tastan, Derya Dilek Kancagi, Bulut Yurtsever, Gamze Tumentemur, Sevda Demir, Raife Dilek Turan, Selen Abanuz, Didem Cakirsoy, Utku Seyis, Samed Ozer, Omer Elibol, Muhammer Elek, Gurcan Ertop, Serap Arbak, Merve Acikel Elmas, Cansu Hemsinlioglu, Ayse Sesin Kocagoz, Ozden Hatirnaz Ng, Sezer Akyoney, Ilayda Sahin, Ugur Ozbek, Dilek Telci, Fikrettin Sahin, Koray Yalcin, Siret Ratip, Ercument Ovali

## Abstract

COVID-19 outbreak caused by SARS-CoV-2 created an unprecedented health crisis since there is no vaccine for this novel virus. Therefore, SARS-CoV-2 vaccines have become crucial for reducing morbidity and mortality. In this study, in vitro and in vivo safety and efficacy analyzes of lyophilized vaccine candidates inactivated by gamma-irradiation were performed. The candidate vaccines in this study were OZG-3861 version 1 (V1), an inactivated SARS-CoV-2 virus vaccine, and SK-01 version 1 (V1), a GM-CSF adjuvant added vaccine. The candidate vaccines were applied intradermally to BALB/c mice to assess toxicity and immunogenicity. Preliminary results in vaccinated mice are reported in this study. Especially, the vaccine models containing GM-CSF caused significant antibody production with neutralization capacity in absence of the antibody-dependent enhancement feature, when considered in terms of T and B cell responses. Another important finding was that the presence of adjuvant was more important in T cell in comparison with B cell response. Vaccinated mice showed T cell response upon restimulation with whole inactivated SARS-CoV-2 or peptide pool. This study shows that the vaccines are effective and leads us to start the challenge test to investigate the gamma-irradiated inactivated vaccine candidates for infective SARS-CoV-2 virus in humanized ACE2+ mice.

## Introduction

Severe acute respiratory syndrome coronavirus 2 (SARS-CoV-2) was first detected in Wuhan, China, in December 2019 and spread globally, causing coronavirus disease 2019 (Covid-19). The number of COVID-19 cases increased at a shocking rate around the world, pushing the limits of “the second wave”. As of 13 December, the total confirmed cases have reached 72,592,974 and the death toll has risen to 1,618,219 (https://www.worldometers.info/coronavirus/). There is still no specific treatment for COVID-19. Several therapies such as various drugs, convalescent plasma, and cellular therapies are under investigation but the efficacy of these treatments is still yet to be improved. In this condition, the urgent need for the SARS-CoV-2 vaccine was responded to by 160 candidates (23 clinical, 137 preclinical) in development ^1^ and some of these candidates reported hopeful results ^2, 3^.

We have previously published our study on the isolation and propagation of the SARS-CoV-2 virus in culture from COVID-19 patients ^4^. In this study, in vitro and in vivo analyzes of our lyophilized vaccine candidates inactivated by gamma-irradiation were performed. Our candidate OZG-3861-01 is a purified inactivated SARS-CoV-2 virus vaccine, and SK-01 is the GM-CSF adjuvant added vaccine candidate. We conducted a preclinical safety and efficacy analysis of the candidates that were applied intradermally to BALB/c mice to assess the toxicity and immunogenicity of OZG-3861-01 and SK-01. Here we report preliminary results including both B cell and T cell response in vaccinated groups. This study leads us to start the challenge test using SARS-CoV-2 viruses and our gamma-irradiated inactivated vaccine candidates in humanized ACE2+ mice.

## Material and Methods

### Sample Collection

Nasopharyngeal and oropharyngeal cavity samples were obtained from four patients who were diagnosed as COVID-19 by Real-Time PCR in Acıbadem Altunizade Hospital, Acıbadem Mehmet Ali Aydınlar University Atakent, and Maslak Hospitals. Informed consent for participation in this study was obtained from participants. In vitro isolation and propagation of SARS-CoV-2 from diagnosed COVID-19 patients were described in our previous study ^4^. The study for SARS-CoV-2 genome sequencing was approved by the Ethics Committee of Acıbadem Mehmet Ali Aydınlar University (ATADEK-2020/05/41) and informed consent from the patients was obtained to publish identifying information/images. These data do not contain any private information of the patients. All techniques had been executed according to the applicable guidelines.

### Manufacturing Gamma-irradiated inactivated SARS-CoV-2 vaccine candidate

For the nasopharyngeal and oropharyngeal swab samples to have clinical significance, it is extremely important to comply with the rules regarding sample selection, taking into the appropriate transfer solution, transportation to the laboratory, and storage under appropriate conditions when necessary ^4^. In **Figure 1**, the production of a candidate vaccine for gamma-irradiated inactivated SARS-CoV-2 was demonstrated. Isolation and propagation were performed from the samples taken on the 7th day when the viral load was predicted to be the most in patients diagnosed with COVID-19. During virus replication, 90% confluent Vero cells in cell culture flasks with a larger surface area were gradually cultured with virus-containing supernatant. The supernatants obtained at the end of the production were pooled and concentrated 10-15 times. To remove cellular wastes in the supernatant, diafiltration was performed. Finally, the concentrated virus was frozen before 50 kGy gamma-irradiation processes. Two different formulations with or without 25ng/ml GM-CSF (CELLGENIX rhGM-CSF) as adjuvants were prepared by the lyophilization stage. Thus, the end products were made available for pre-clinical in vitro and in vivo analyzes.

**Figure 1:**
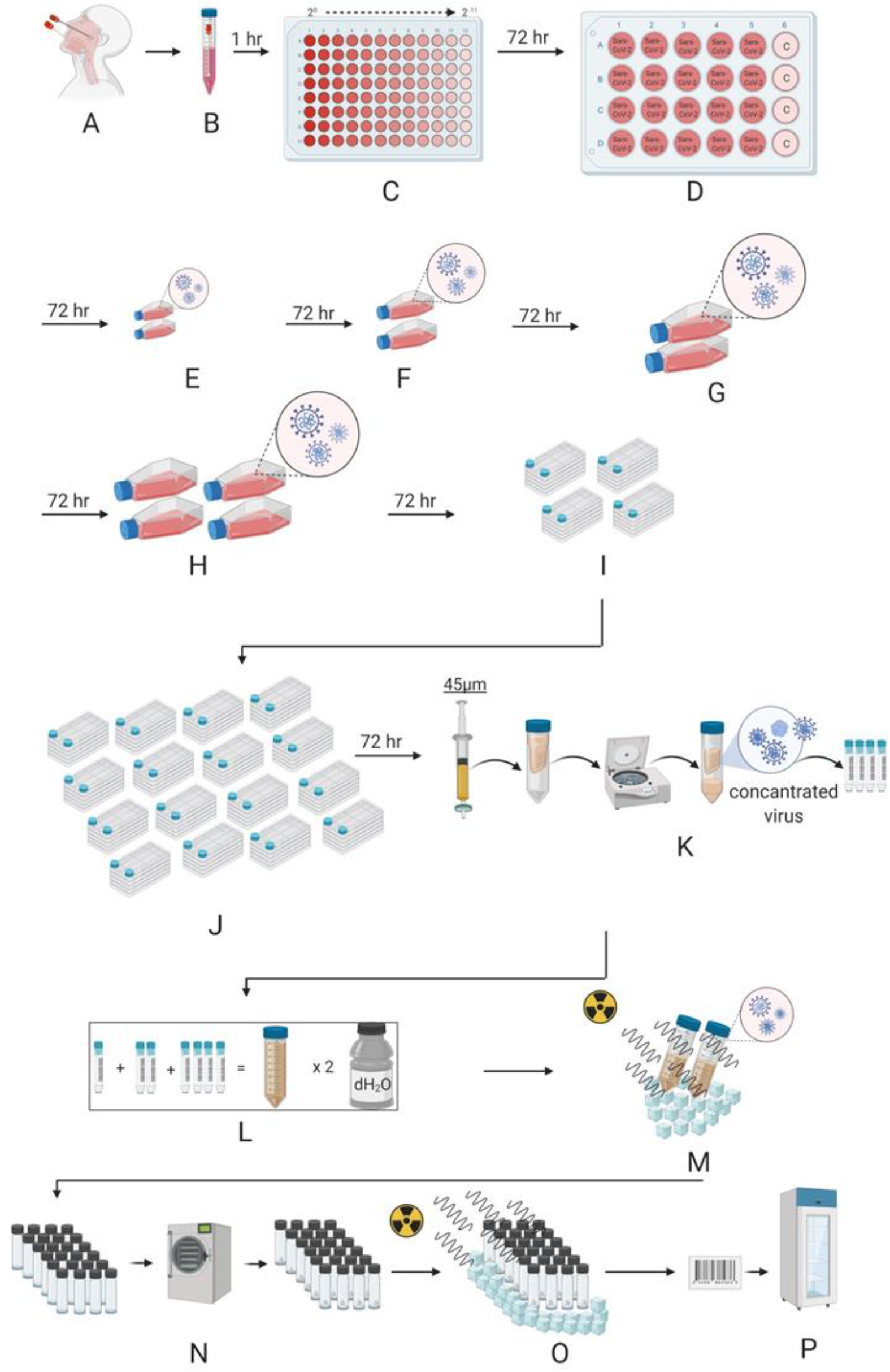
Representation of Gamma-irradiated inactive lyophilized SARS-CoV-2 manufacturing. **A.** Nasopharyngeal and Oropharyngeal samples were taken. **B.** Sample came to the laboratory in a 2-8 C transfer solution. **C.** The virus was distributed by making serial dilution (up to 2^-11^) onto Vero cells. Viruses were transferred to **D.** a 24-well plate **E.** T-75 flasks F. T-175 flasks **G.** and **H.** T-300 flasks with a confluent with Vero cells by increasing culturing surface area. Next, the propagated virus was transferred to **I.** four and **J.** sixteen multi-layered flasks with a confluent with Vero cells. **K.** The total virus solution was then passed through a 45 μm filter, the virus was concentrated by centrifugation in a special tube with a 100 KDa filter. The concentrated virus was stored at −80 C before irradiation. **L.** All concentrated viruses obtained are pooled and washed two times with distilled water for diafiltration in a 100 KDa concentrator. **M.** The concentrated virus mixture was inactivated by irradiation at 25 kGy in dry ice. **N.** Inactivated virus is lyophilized after dose adjustment. **O.** The lyophilized virus mixture is sterilized by irradiation at 25 kGy in dry ice. **P.** The lyophilized bottled inactive SARS-CoV-2 vaccine is labeled and stored at 4 C.

### Viral RNA Extraction and Viral Genome Sequencing

Viral RNA extractions were performed by QUICK-RNA Viral Kit (Zymo Research, USA) in the Acıbadem Labcell Cellular Therapy Laboratory BSL-3 Unit according to the manufacturer's protocols. Library preparation was performed by CLEANPLEX SARS-CoV-2 Research and Surveillance NGS Panel (Paragon Genomics, USA) according to the manufacturer’s user guide. For the construction of the library, The CLEANPLEX Dual-Indexed PCR Primers for ILLUMINA (Paragon Genomics, USA) were used by combining i5 and i7 primers. Samples were sequenced by ILLUMINA MiSeq instrument with paired-end 131 bp long fragments. The data that passed the quality control were aligned to the reference genome (NC_045512.2) in Wuhan and a variant list was created with variant calling. The data analysis was described in detail in our previous study ^5^.

### NANOSIGHT

Nanoparticle Tracking Analysis (NTA) measurements were carried out for SARS-CoV-2 titer in suspension by using The NANOSIGHT NS300 (Amesbury, UK). Samples were diluted with distilled water at a 1:10 ratio and transferred to NANOSIGHT cuvette as 1 ml. Measurements were performed at room temperature with 5 different 60-second video recording.

### ZETA analyzing

Dynamic light scattering (DSL) measurements of SARS-CoV-2 were carried out using a ZETASIZER nano-ZS from Instruments (Malvern, UK). Samples were diluted with distilled water 1:10 ratio and transferred to a polystyrene cuvette (10 mm). The volume of the analyzed preparations was 1 ml. Measurements were performed at room temperature with a He-Ne laser (633 nm, 10 mW) and scattered light detection at 173°. Measured data were processed using the Dispersion Technology Software version 5.10.

### RT-PCR

Total RNA isolations from SARS-CoV-2 were carried using DIRECT-ZOL RNA Miniprep Kits (Zymo Research, USA), and concentrations were determined using QUBIT fluorometer with the QUBIT RNA HS Assay (Thermo Fisher Scientific, USA). SARS-CoV-2 specific RT-PCR was performed with BOSPHORE Novel Coronavirus (2019-nCoV) Detection Kit (Anatolia Geneworks, Istanbul) along with Orf1ab and E gene primers. The RT-PCR analysis was performed in ROCHE Lightcycler 96.

### Quantitative RT-PCR to determine viral copy number

Total RNA isolations were performed from SARS-CoV-2 specimens using DIRECT-ZOL RNA Miniprep Kits (Zymo Research, USA). Quantitative RT-PCR was performed with the QUANTIVIRUS SARS-CoV-2 Test Kit (Diacarta) according to the manufacturer’s protocol. The quantitative RT-PCR analysis was analyzed in ROCHE LIGHTCYCLER 96.

### Inactivated SARS-CoV-2 virus imaging by transmission electron microscopy

Viruses were inactivated and fixed with 2.5% glutaraldehyde in PBS (0.1 M, pH 7.2) for 2.5 h. One drop of glutaraldehyde-treated virus suspension was placed on the carbon-coated grid for 10 min. The remaining solution was absorbed with a filter paper and the grid was stained by a negative staining procedure. Then, it was evaluated under a transmission electron microscope (Thermo Fisher Scientific- TALOS L120C) and photographed.

### LC-MSMS Protein Analysis

LC-MSMS protein analysis was performed at Acibadem Labmed Laboratory, Istanbul. ONAR data acquisition mode was applied by a WATERS XEVO G2-XS high-resolution mass spectrometer. Tryptic peptides were generated by overnight digestion with trypsin followed by reduction and alkylation steps with DTT and IAA, respectively, and fractionated by a 90 min reverse-phase gradient at 500 nL/min flow rate on an HSS T3 (Waters-186008818) nano column. LC-MSMS data was searched against the NCBI RefSeq sequence database for protein identification. PROGENESIS QIP software was used for protein identification (Waters v4.1).

### Replicative Competent Coronavirus test with gamma-irradiated inactivated SARS-CoV-2 vaccine candidates

3μg of lyophilized inactivated SARS-CoV-2 vaccine candidate in 100 μl apyrogenic water was inoculated into %90 confluent Vero cells at 37C. The supernatant of this culture was replenished with fresh Vero cell culture every 3-to-5 days up to 21 days of incubation. As a negative control, only 100 μl apyrogenic water was inoculated into Vero cells and cultured for 21 days with the same treatments. At the end of the incubation, the final supernatant was collected, centrifuged at 2000G for 10 min to remove cell debris. Next, the supernatants were concentrated 10x with 100kDa Amplicon tubes. The concentrated samples were tested in the XCELLIGENCE RTCA system in a dose-dependent manner as 10^-1^ to 10^-6^ to determine the cytopathic effect.

### BALB/c mice

For studies on BALB/c mice, we confirm that all methods were carried out in accordance with relevant guidelines and regulations. Furthermore, we confirmed that the study was carried out in compliance with the ARRIVE guidelines. Female or male 11-months-old BALB/c mice were housed in AAALAC International accredited Acıbadem Mehmet Ali Aydinlar University Laboratory Animal Application and Research Center (ACUDEHAM; Istanbul, Turkey) for 7-day toxicity and 21-day toxicity and efficacy tests. Light, temperature, humidity, and feeding conditions followed the ACUDEHAM accredited operating procedures. For 34-day efficacy tests, female or male 3-months-old BALB/c mice were housed in Yeditepe University Experimental Research Center. All animal studies received ethical approval by the Yeditepe University Animal Experiments Local Ethics Committee (Yeditepe-HADYEK). Mice in the intervention groups were identified as female and male plus a correlative number 1–10. Cages were identified with the study name and color codes. Each mouse was marked with its code including vaccine treatment with/without adjuvant in the base of the tail using a non-toxic permanent marker as single, double, or triple line and without a line.

### In Vivo Inactivated Vaccine Candidate Treatments

To determine the 21-day immunogenicity (n=3/group) and 7-day (n=4/group) or 21-day toxicity (n=3/group) of inactive vaccine produced in Acibadem Labcell Cellular Therapy Laboratory, Istanbul, Turkey, on day 0 mice were inoculated with the dose of 3 μg/100 μl (4,2×10^6^ SARS-CoV-2 viral copy per microgram) adjuvanted or nonadjuvanted vaccine intradermally and with apirogen water in the control group. In two other groups, a booster dose of 3 μg/100 μl adjuvanted or nonadjuvanted vaccine was administered on day 15 intradermally in addition to day 0. Survival and weight change were evaluated daily and every week respectively. To evaluate the fast response toxicity, on day 0 mouse was inoculated with the dose of 3 μg/100 μl adjuvanted or nonadjuvanted vaccine intradermally and with apirogen water in the control group (n=4/group). Survival and weight change were evaluated on days 0, 3, and 7. Blood samples were collected just before the sacrification for hemogram and biochemical analysis on day 7. For long term toxicity and immunogenicity, blood samples were collected just before the sacrification on day 21 or day 34 for serum preparation to be used in preclinical in vitro studies. Mice treated with the vaccine candidates were sacrificed on day 21 or day 34 postimmunization for analysis of B and T cell immune responses via SARS-Cov-2 specific IgG ELISA, IFNγ ELISPOT, and cytokine bead array analysis.

### Histopathological Applications

Mice treated for both toxicity and efficacy tests were sacrificed on day 7 or day 21 postimmunization for histopathology analysis. Dissected organs including the cerebellum, lungs, liver, kidneys, skin, intestine, and part of the spleen of sacrificed mice were taken into 10% buffered formalin solution prior to routine tissue processing for histopathological analysis. Tissue tracking was performed firstly in NBF 10% for 1 hour and then in alcohol from 60% to absolute gradually for 1 hour/each alcohol concentration. The tissue tracking was finalized in Xylene and Paraffin for 1 hour/each. Blocking of tissues was performed by embedding them in paraffin and turned into blocks. Sections with 3-4 μm thickness were taken from paraffin blocks. Next, staining was performed following several procedures including deparaffinization, hydration, hematoxylin stage, acid alcohol phase, bluing, eosin phase, dehydration, transparency step, and closing with the non-aqueous closing agent.

### SARS-CoV-2 IgG ELISA

Prior to the sacrification, blood samples were collected from the whole group of mice. The serum was collected with centrifugation methods. Serum samples were stored at −40 C. To detect the SARS-COV-2 IgG antibody in mouse serum SARS-COV-2 IgG ELISA Kit (Creative, DEIASL019) was used. Before starting the experiment with the whole sample, reagent and microplates pre-coated with whole SARS-CoV-2 lysate were brought to room temperature. As a positive control, 100 ng mouse SARS-CoV-2 Spike S1 monoclonal antibody was used which is commercially available (E-AB-V1005, Elabscience). Serum samples were diluted at 1:64, 1:128, and 1:256 in a sample diluent, provided in the kit. Anti-mouse IgG conjugated with Horseradish peroxidase enzyme (mHRP enzyme) was used as a detector. After incubation with the stoping solution, the color change was read at 450nm with the microplate reader (OMEGA ELISA Reader).

### Neutralization assay using Real-Time Cell Analysis (RTCA), XCELLIGENCE

TCID50 (Median Tissue Culture Infectious Dose) of SARS-CoV-2 was determined by incubating the virus in a serial dilution manner with the Vero cell line (CCL81, ATCC) in gold microelectrodes embedded microtiter wells in XCELLIGENCE Real-Time Cell Analysis (RTCA) instruments (ACEA, Roche) for 8 days. Neutralization assay was performed with 1:64, 1:128, and 1:256 dilutions of mice serum pre-incubated with a 10X TCID50 dose of SARS-CoV-2 at room temperature for 60 min. Infective active SARS-CoV-2 virus to be used in neutralization tests was titrated in the RTCA system and the dose of TCID50 was determined. It was decided to use 10 times more than the dose of TCID50 in the following neutralization tests as 100X TCID50 dose. Next, the pre-incubated mixture was inoculated into the Vero-coated cells which were analyzed in real-time for 120 hours (totally, 145 hours). Cell analysis was normalized to the value at the 24th hour of culturing before culturing with serum-SARS-CoV-2 sample conditions. Normalized cell index shows the proliferation and viability of the adherent cells (the higher cell index means the higher viability and proliferation). The neutralization ratio was determined by assessing percent neutralization by dividing the index value of serum-virus treated condition wells by the cell index value of untreated control Vero cells (normalized to 100%). For example, for the sample of 1:128 adjuvant+ double-dose, the normalized cell index value was 0,651 while the index value of the control well was 0,715. At this time point, the cell index value of only virus incubated wells was 0. This gave 91,4% virus neutralization. This calculation was performed for each mouse in the group and the mean of the virus neutralization was determined.

### Antibody-Dependent Enhancement Assay using qRT-PCR

Peripheral blood mononuclear cells (PBMC) from healthy donor blood was isolated using the Ficoll-Paque solution. PBMCs were cultured in the T-300 flask for 2 hours at 37 C. Non-binding cells (T cells) were discarded by withdrawing the medium after the incubation. Following washing, flask-attached cells were mostly monocytes that were cultured in XCELLEGENCE plates for 24 hr before incubation with mice serum and SARS-CoV-2. A mice serum dose of 1:256 was preincubated with a dose of 100x TCID50 SARS-CoV-2. After 48 hours of incubation on the monocytes, qRT-PCR was performed by scraping off the supernatant and cells to assess the SARS-CoV-2 copy number per ml.

### Cytokine Bead Array (CBA) From Serum

MACSPLEX Cytokine 10 kit (Miltenyi Biotec) was used for the Cytokine bead array following the manufacturer’s protocol. To study the CBA test from serum samples, serum samples were diluted 1: 4 and tested. Samples were collected into sample tubes, and flow analysis was done. Flow analysis was performed with the MACSQUANT Analyzer (Miltenyi Biotec).

### Mouse IFN-γ ELISPOT analysis

Mouse Spleen T cells were centrifuged with Phosphate Buffer Saline (PBS) at 300xg for 10 min. Pellet was resuspended in TEXMACS (Miltenyi Biotech, GmbH, Bergisch Gladbach, Germany) cell culture media (%3 human AB serum and 1% Pen/Strep). 500,000 cells in 100 μl were added into microplate already coated with a monoclonal antibody specific for mouse IFN-γ. Either 3μg/ 100 μl inactivated SARS-CoV-2 or 1000 nM SARS-CoV-2 virus PEPTIVATOR pool (SARS-CoV-2 S, N, and M protein peptide pool) (Miltenyi Biotech, GmbH, Bergisch Gladbach, Germany) were added into each well including mouse spleen T cells. The microplate was incubated in a humidified 37°C CO_2_ incubator. After 48-72 h incubation, IFN-γ secreting cells were determined with Mouse IFNγ ELISPOT Kit (RnDSystems, USA) according to the manufacturer’s instructions. The spots were counted under the dissection microscope (Zeiss, Germany).

### Stimulated T cell cytokine response and immunophenotype

500,000 cells isolated from mouse spleen were incubated with 1000 nM SARS-CoV-2 virus PEPTIVATOR pool (SARS-CoV-2 S, N, and M protein peptide pool) (Miltenyi Biotech, GmbH, Bergisch Gladbach, Germany) in a humidified 37°C CO_2_ incubator. After 48-72 h incubation, the mouse cytokine profile was analyzed using the supernatant of the cultures using the MACSPLEX Cytokine 10 kit (Miltenyi Biotec). Also, in order to determine T cell activation and proliferation, the restimulated cells were stained with the antibodies including CD3, CD4, CD8, CD19, and CD25 as an activation marker (Miltenyi Biotec). The Cytokine bead array and the T cell activation and proportions were analyzed using the MACSQUANT Analyzer (Miltenyi Biotec).

### Statistics

Normally distributed data in bar graphs was tested using student’s t-tests for two independent means. The Mann-Whitney U test was employed for comparison between two groups of non-normally distributed data. Statistical analysis of the presence or absence of toxicity including inflammation in the tissue sections was performed using the Chi-squared test. Statistical analyses were performed using SPSS Statistics software. No outliers were excluded in any of the statistical tests and each data point represents an independent measurement. Bar plots report the mean and standard deviation of the mean. The threshold of significance for all tests was set at *p<0.05. *NS* is Non-Significant.

## Results

### Manufacturing Gamma-irradiated inactivated SARS-CoV-2 vaccine candidate

Most of the therapeutic options available to treat COVID-19 are based on previous experience in the treatment of SARS- and MERS-CoV ^6^. The main reason for the lack of approved and commercially available vaccines or therapeutic agents against these CoVs may be the relatively high cost and long production time ^6^. Multiple strategies have been adopted in the development of CoV vaccines; most of these are recombinant adenovirus-based vaccines and immuno-informatics approaches used to identify cytotoxic T lymphocyte (CTL) and B cell epitopes ^7, 8^. Unlike the vaccine obtained with the recombinant protein cocktail of the virus, the whole of the virus in the vaccine candidates may enable to produce a vast amount of the immunoglobulin molecules that can recognize the virus antigens better and more specifically. With our straightforward manufacturing protocol of the whole inactivated lyophilized SARS-CoV-2 vaccine, two different formulations with or without GM-CSF as adjuvants were prepared (**Fig. 1**). Furthermore, since the inactivated virus vaccine manufacturing process would require careful characterization of viral isolates as seeds, and demonstration of consistent in viral cultures, we have shown that our inactivated virus-based vaccine production procedure meets the criteria with the following three independent vaccine production (**Supplementary Table 1**). Thus, the end products were made available for pre-clinical in vitro and in vivo safety and efficacy analyzes.

### Genome Sequencing of the SARS-CoV-2 isolates

While evaluating the appropriate isolate for the inactive vaccine form, viral genome sequencing obtained from four patients was performed and a variant list was created (**Table 1**). Representative IGV reads from each patient were depicted in **Figure 2**. The variants detected in patients were identified in previous sequencing results as well. Only one variant was novel according to the analysis in the GISAID database. The effect of the variants on the protein level and multiple alignment analysis results were presented in our viral genome sequencing study ^5^.

**Table 1.** Identified variants in four patients.

| **Nucleotide position** | **Gene/region** | **Gene product** | **Nucleotide exchange (Ref/Alt)** | **Amino acid exchange** | **Mutation type** | **Conservation among 9332 samples** | **Frequency in this study** | **Detection in previous studies** |
| --- | --- | --- | --- | --- | --- | --- | --- | --- |
| **22227** | S | surface glycoprotein | C/T | A222V | missense | 99.09% | 15.4% | Novel |
| **241** | 5' UTR | non-coding | C/T | - | non-coding | 59.52% | 53.8% | Detected |
| **2113** | ORF1ab | Nsp2 | C/T | I436I | synonymous | 98.07% | 61.5% | Detected |
| **3037** | ORF1ab | Nsp3 | C/T | F106F | synonymous | 61.27% | 30.8% | Detected |
| **7765** | ORF1ab | Nsp3 | C/T | S1682S | synonymous | 98.18% | 15.4% | Detected |
| **14408** | ORF1ab | RNA-dependent RNA Polymerase | C/T | P323L | missense | 61.01% | 23.1% | Detected |
| **17523** | ORF1ab | Helicase | G/T | M429I | missense | 98.50% | 23.1% | Detected |
| **17690** | ORF1ab | Helicase | C/T | S485L | missense | 98.02% | 61.5% | Detected |
| **18877** | ORF1ab | 3'-to-5' exonuclease | C/T | L280L | synonymous | 96.13% | 30.8% | Detected |
| **23403** | S | Surface glycoprotein | A/G | D614G | missense | 61.39% | 15.4% | Detected |
| **25563** | ORF3a | ORF3a protein | G/T | Q57H | missense | 71.27% | 53.8% | Detected |

**Figure 2:**
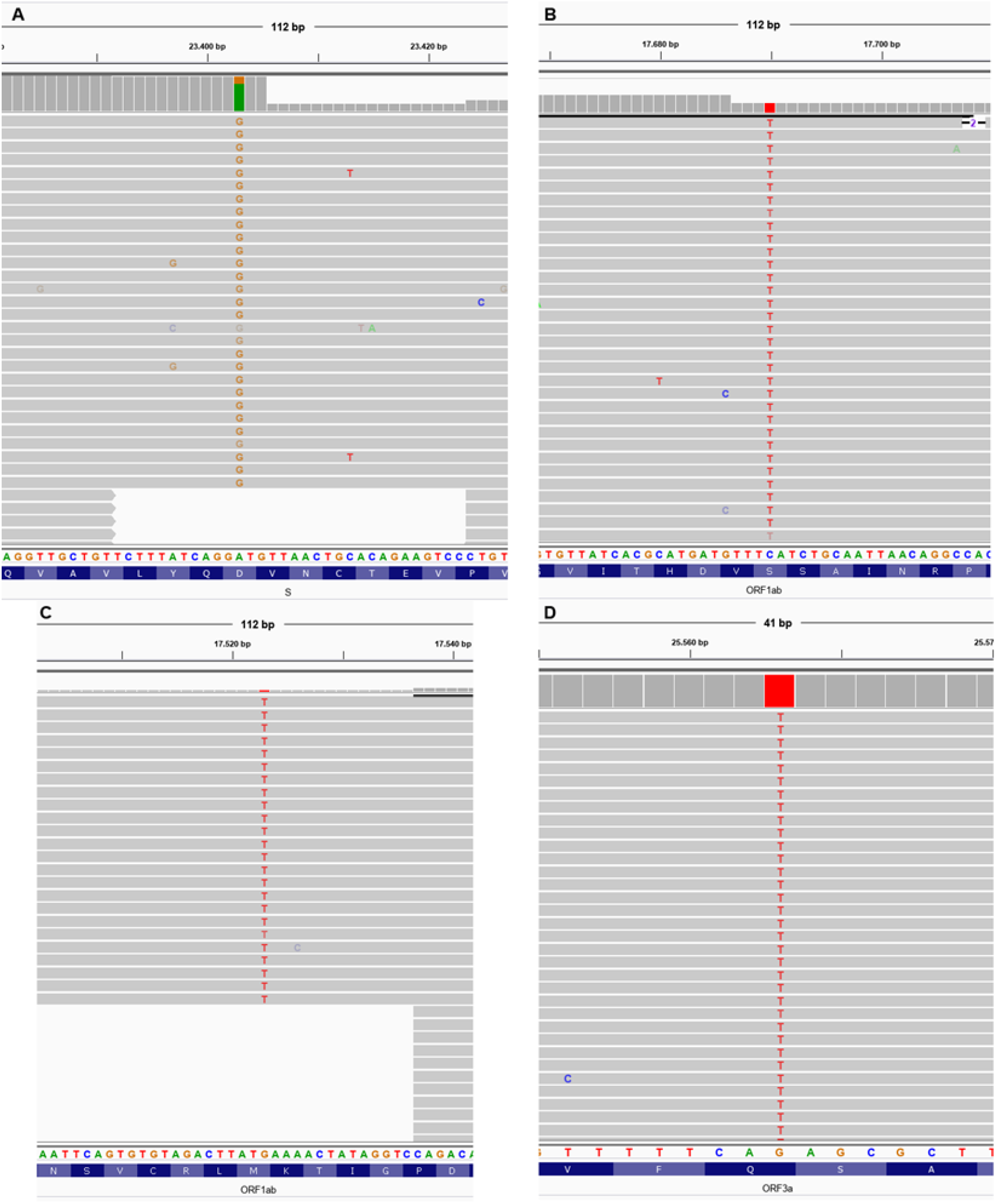
Representative IGV imaging of detected variants. **A.** ACUTG-1, D614G missense SNV (A23403G) on the surface glycoprotein. **B.** ACUTG-2, S485L missense SNV (C17690T) on helicase. **C.** ACUTG-3, M429I missense SNV (G17523T) on helicase. **D.** ACUTG-4, Q57H missense SNV (G25563T) on ORF3a protein.

### Characterization and quantification of final product gamma-irradiated inactivated SARS-Cov-2

Pooling was performed to obtain the final product with SARS-CoV-2 isolates which were sequenced genome. RT-PCR identification of the isolates was performed as stated in our previous publication ^4^. The dry end product obtained after inactivation and lyophilization by gamma-irradiation was diluted to 3 ug / 200 μl and analyzed to measure particle count, size, and density. As a result of these analyzes, the average size of the particles in SK-01 V1 was 139.3 _+/−_ 5.6 nm (**Fig. 3A and 3B**) and the average size of the particles in OZG-3861 V1 was determined to be 144 nm _+/−_ 51.8 nm (**Fig. 3C and 3D**). However, the particle density in this size range was calculated to be 91.9% _+/−_ 2.5% (**Fig. 3D**). The number of viral particles per dose was found to be 2.6×10^8^ _+/−_ 2.61×10^7^. Results illustrate that the virus particles in the final product largely retain their compact structure. However, negative staining and transmission were analyzed with an electron microscope to display the compact structure of the virus particles in the final product (**Fig. 4A**). In addition to RT-PCR analysis, the presence of SARS-CoV-2 specific protein sequences including Replicase polyprotein 1ab and Non-structural protein 3b) was confirmed by proteome analysis on the final product, **figure 4B** shows eluted peptides between m/z 50-2000 along a 90 min reverse-phase gradient elution. At the same time, the gamma-irradiated inactive virus strains lost their infective properties was confirmed by the replicative competitive coronavirus test using the RTCA analysis performed at the end of the 21-day repeated passage (**Fig. 4C**). Also, to demonstrate the reliability and consistency of the inactivation assay, we repeated the test with colorimetric MTT assay post 21-day culturing of the inactivated virus samples. We showed inactivated virus samples in three representatives of inactivated vaccine samples (**Supplementary figure 1**). As a result of these analyzes, it has been decided that vaccine candidates have been made final products for use in toxicity and efficacy studies.

**Figure 3:**
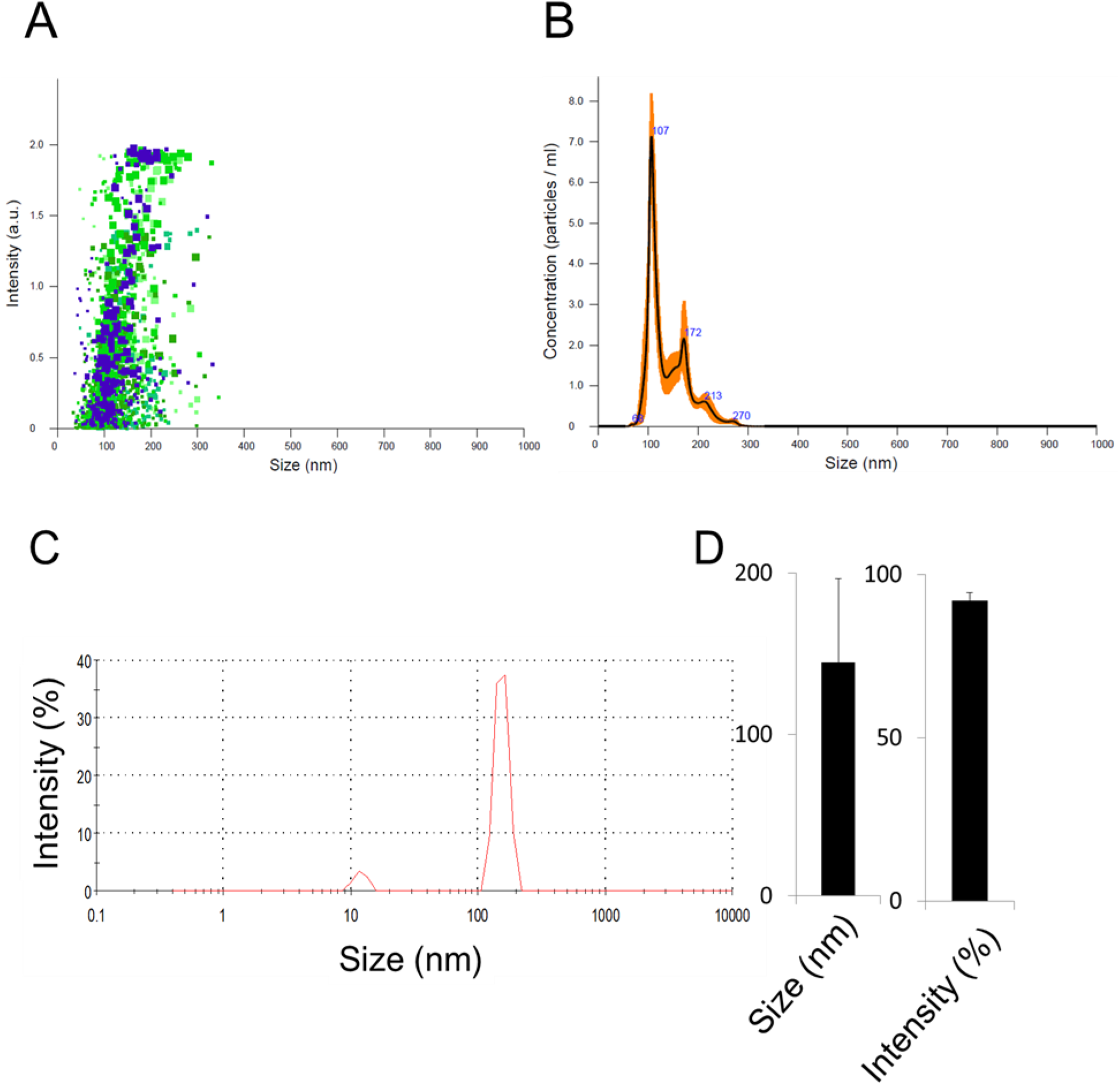
Quantification of particle number, size, and intensity in lyophilized SARS-CoV-2. **A.** Plot showing intensity versus the size of the particles in SK-01 V1 (inactivated virus & GM-CSF). **B.** Plot showing means of particle size of SK-01 V1 in the sample read three times concerning the concentration. **C.** Graph illustrating zeta analysis of inactivated virus particles in OZG-3861 V1 concerning intensity. **D.** Bar graphs showing quantified size and intensity of the sample.

**Figure 4:**
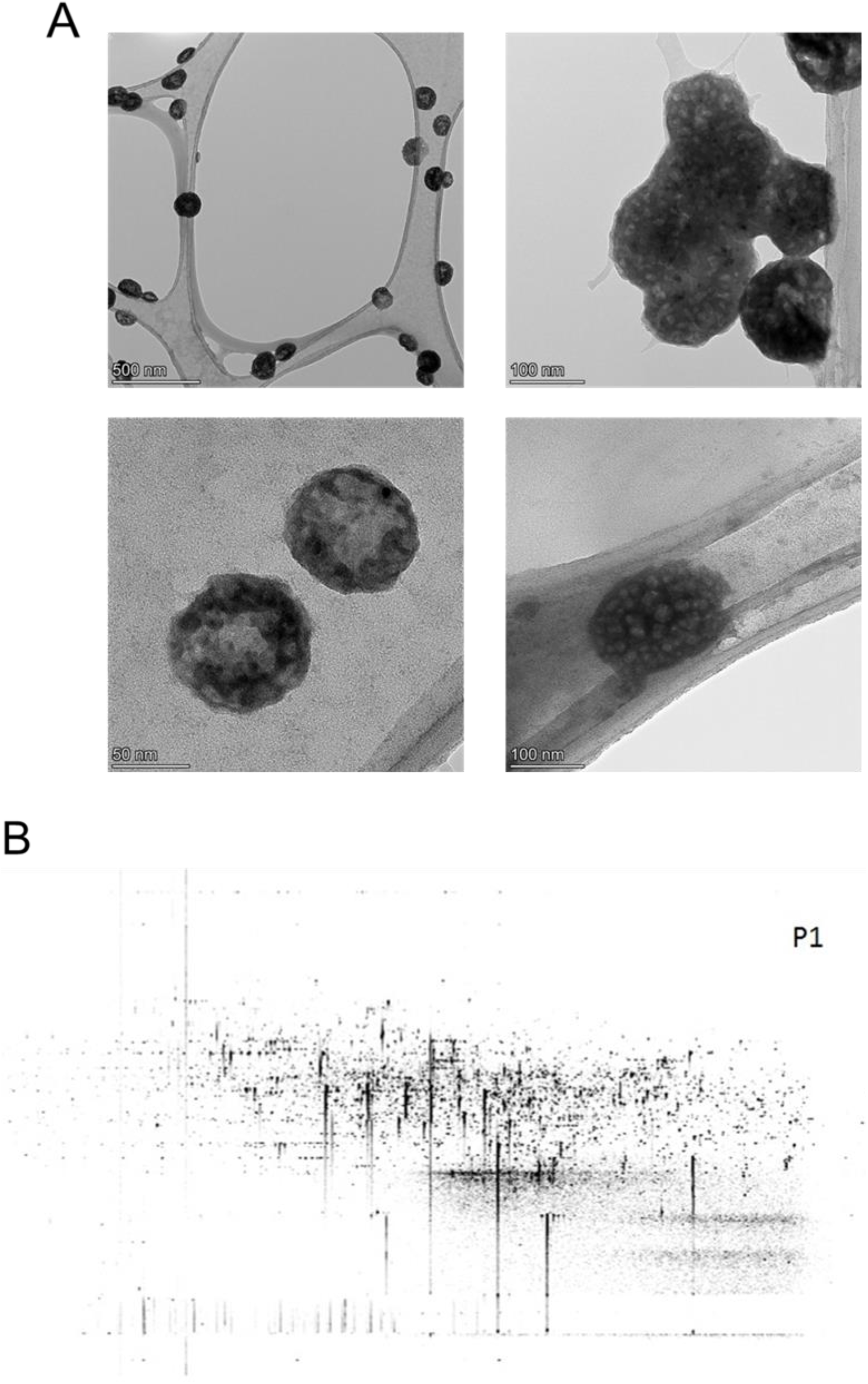
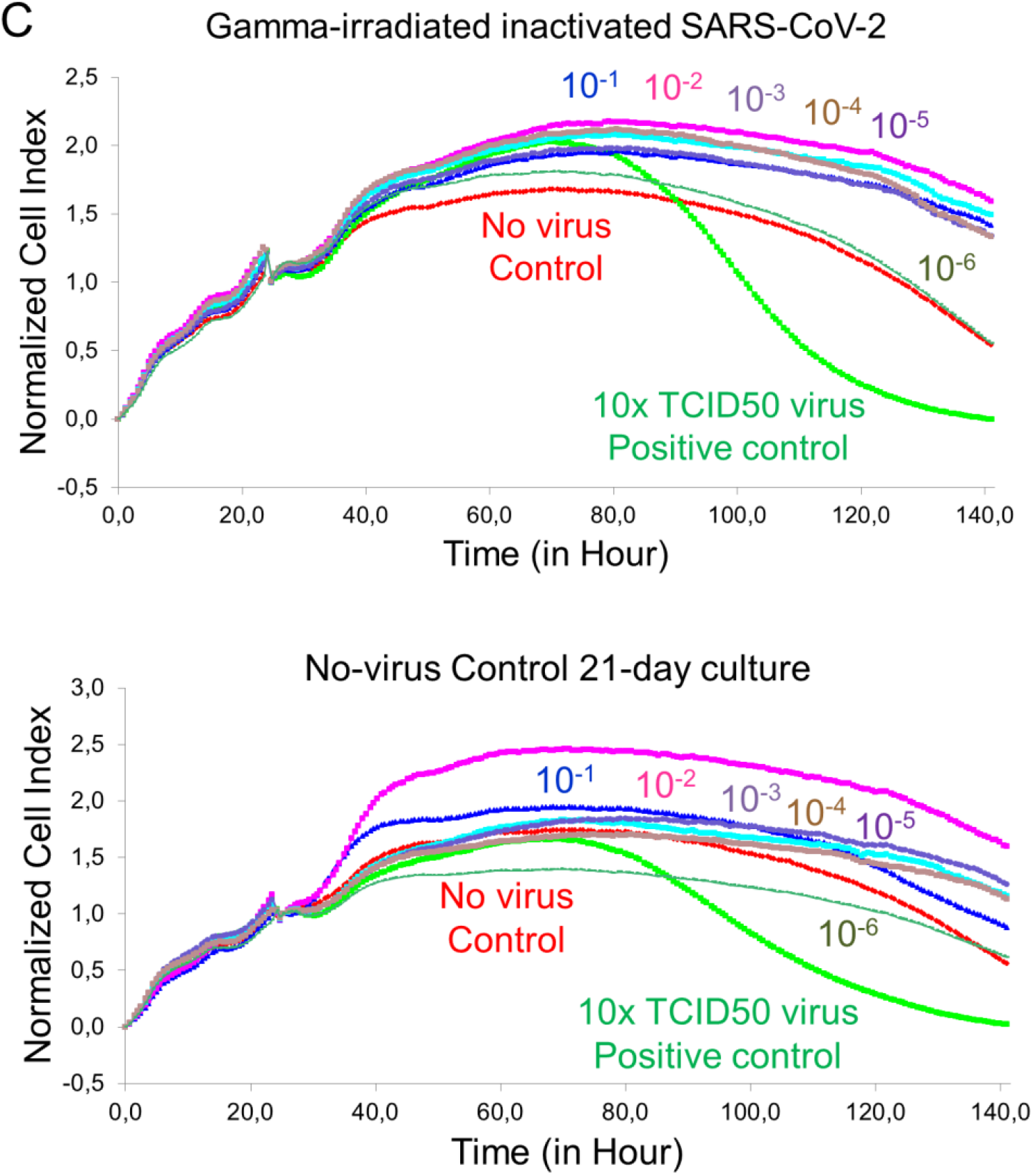
Transmission Electron Microscopy Imaging and Proteome Analysis. **A.** Representative electron micrographs of inactivated SARS-CoV-2. The group of virus particles was seen on the grid (Scale bars: 500 nm, 100 nm, 50 nm). **B.** Proteome analysis of inactivated SARS-CoV-2 product. **C. G**raphs showing real-time cell analysis of gamma-irradiated inactivated SARS-CoV-2 and no—virus control cultured on Vero for 21 days to determine the presence/absence of cytopathic effect in a dose-dependent manner. The Red line is no virus Vero internal control and the Green line is 10x TCID50 doses of infective SARS-CoV-2 as a positive control.

### Safety analysis of the vaccine candidates, SK-01 V1 and OZG-3861 V1

In order to test the reliability including the 7-day and 21-day toxicity of the vaccine candidates, intradermal administration was performed to the mouse groups as a single dose with adjuvant (SK-01 V1) and single-dose without adjuvant (OZG-3861 V1) (**Fig. 5A**). At a one-week follow-up, no significant weight change was detected in groups compared to the control mouse group (**Fig. 5B**). There was also no significant difference in CBC analysis (red blood cell, RBC; white blood cell, WBC; hemoglobin; HGB and platelet rates) (**Fig. 5C and 5D**). However, when the study groups were compared with the control, there was a significant increase in gammaglobulin and related protein increase in the vaccine group containing adjuvant (**Fig. 5E**). In toxicity analyzes, Ca, ALT, and LDH values did not differ significantly between the groups (**Fig. 5F, 5G, and 5H**). In the histopathological analysis on day 7, no difference was observed in the samples of spleen, liver, lung, intestine, hippocampus, kidney, and skin among the groups (**Fig. 5I**). In the examination of cerebellum tissues, no statistically significant pathology in comparison with the control group was observed (**Fig. 5I**). In the adjuvant negative single dose (OZG-3861 V1) group, dense Purkinje cells were observed. However, this density did not appear to be significant (p >0.05) in comparison with the control group (**Fig. 5I**). These toxicity analyzes encouraged us to start an *in vivo* efficacy and dose study with both adjuvant SK-01 V1 and OZG-3861 V1 vaccine candidates without adjuvant in mice.

**Figure 5:**
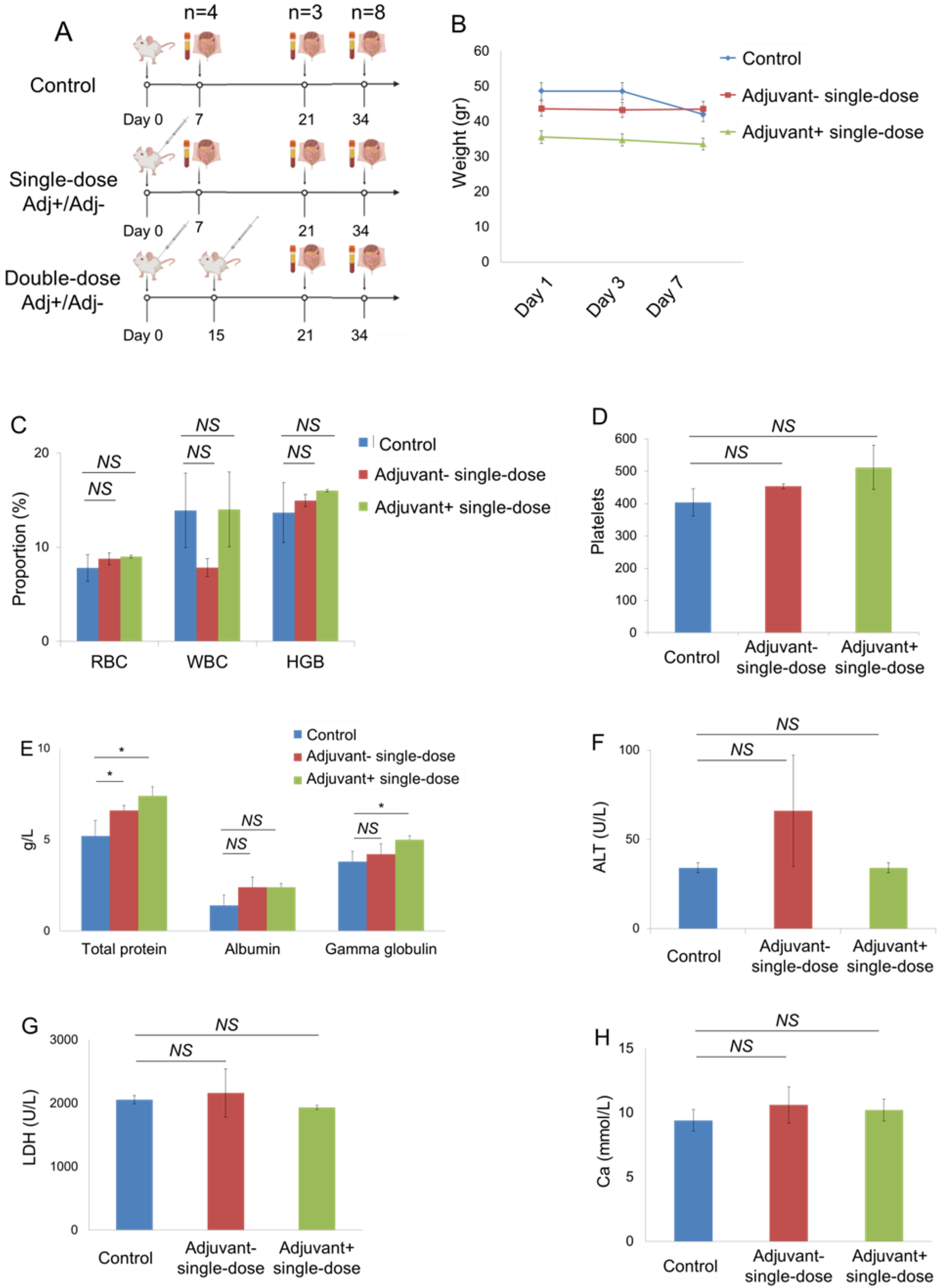
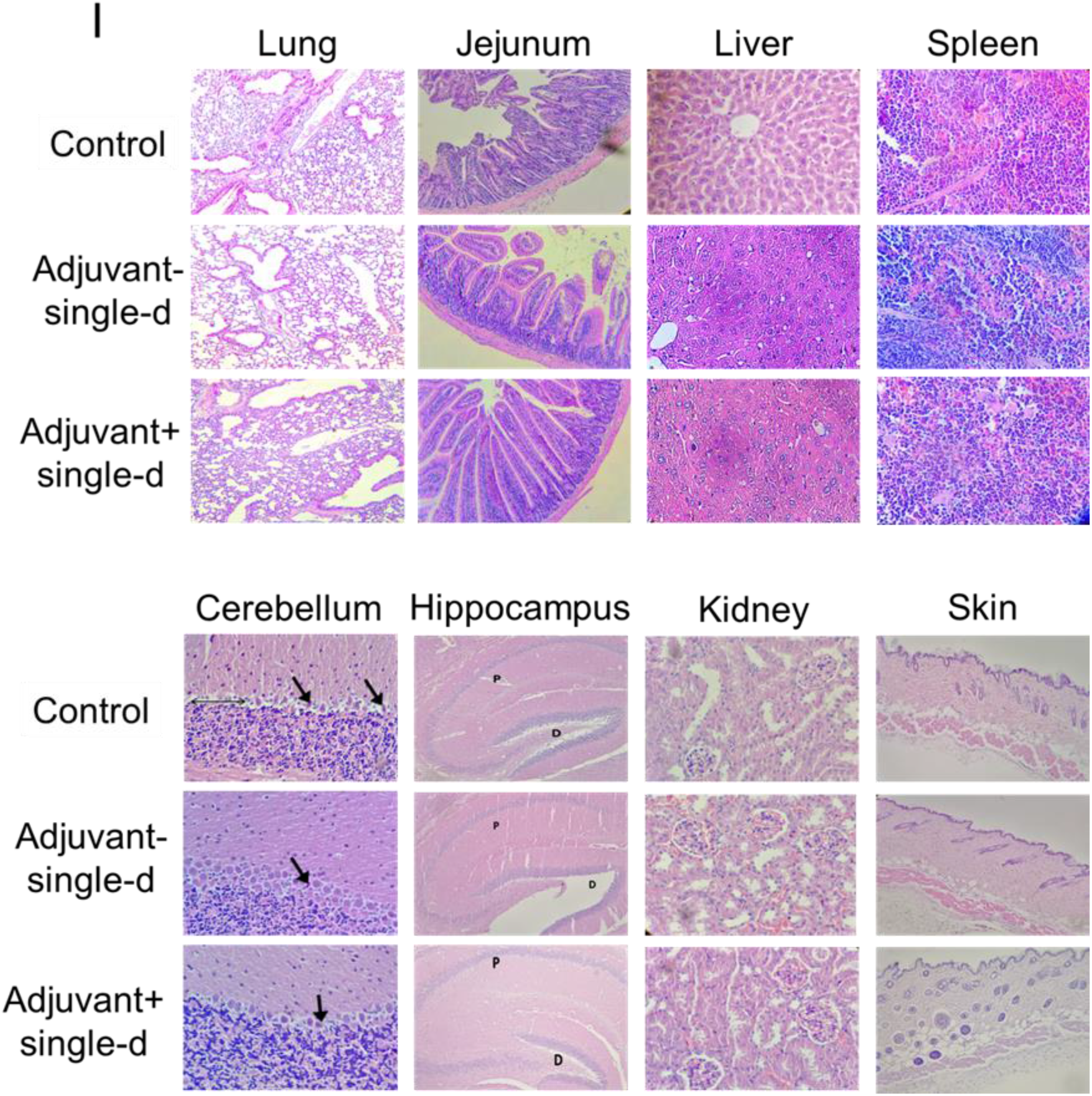
Day-7 Safety analysis of the vaccine candidates, SK-01 V1, and OZG-3861 V1. **A.** In vivo inactivated vaccine candidate treatments and euthanizations. Experimental plan of in vivo SK-01 V1 and OZG-3861 V1 intradermal treatment as single or double dose. **B.** Graph showing weight change during one week in groups; control, blue; adjuvant negative single-dose (OZG-3861 V1), red; adjuvant positive single-dose (SK-01 V1), green. **C.** Bar graph showing hemogram analysis including RBC, WBC, and HGB levels in the groups. **D.** Bar graph showing the change in platelet proportions between groups in one week. **E.** Bar graph showing levels of total blood protein, albumin, and gamma-globulin (g/L) in the groups. Bar graphs showing levels of **F.** ALT (U/L), **G.** LDH (U/L) and **H.** Ca (mmol/L) in the blood of mice groups. **I.** Histopathologic analysis on day 7 of the lung, jejunum of intestine, liver, spleen, cerebellum, hippocampus, kidney, and skin. Purkinje neurons (arrow). The thin double-headed arrow is space, the thick arrow is a picnotic cell.

Subsequently, long-term toxicity analysis at day 21 was determined. Histopathology analysis showed no significant pathological finding in the lung, liver, jejunum of intestine, spleen, cerebellum, hippocampus, kidney, and skin tissues (**Fig. 6A & 6B**). Numerous foci of megakaryocytes (marked by a star) and trabeculae (marked by arrow) were determined in the histological sections of the spleen in all groups (**Fig. 6A**). Cerebellum sections were studied in brain tissues obtained from mouse groups. In particular, there was no statistically significant difference in the shape and staining properties of Purkinje neurons in the cerebellum cortex of all groups. Interestingly, the cells in the adjuvant-negative single-dose group had better shapes in comparison with the other groups (**Fig. 6B**). No pathological finding was observed in the dentate gyrus and the pyramidal layer of the hippocampus in all groups (**Fig. 6B**). Distal and proximal tubules in the kidney were observed similarly in all groups (**Fig. 6B**). On the other hand, a statistically significant (p <0.05) inflammatory reaction was observed in the analyzed skin and kidney tissues in the adjuvant positive double dose vaccine administration group (**Fig. 6B**). Furthermore, glomerulus structures in all vaccinated groups were normal. In toxicity analysis of skin tissue, inflammatory cells infiltration, and eosinophils in some dermis area of the vaccination points of the skin were detected in the double-dose groups (**Fig. 6B**). The vaccine candidates had no significant toxicity on the tissues. The analysis was also performed to investigate whether there was an increase in Th1, Th2, and Th17 dependent cytokine releases in the blood sera collected from the mouse groups that received the vaccine either at day 21. Compared to the control groups, no statistically significant cytokine increase was observed in any application group (**Fig. 6C**). Findings show that there was no toxic side effect of SK-01 V1 and OZG-3861 V1 in mouse groups.

**Figure 6:**
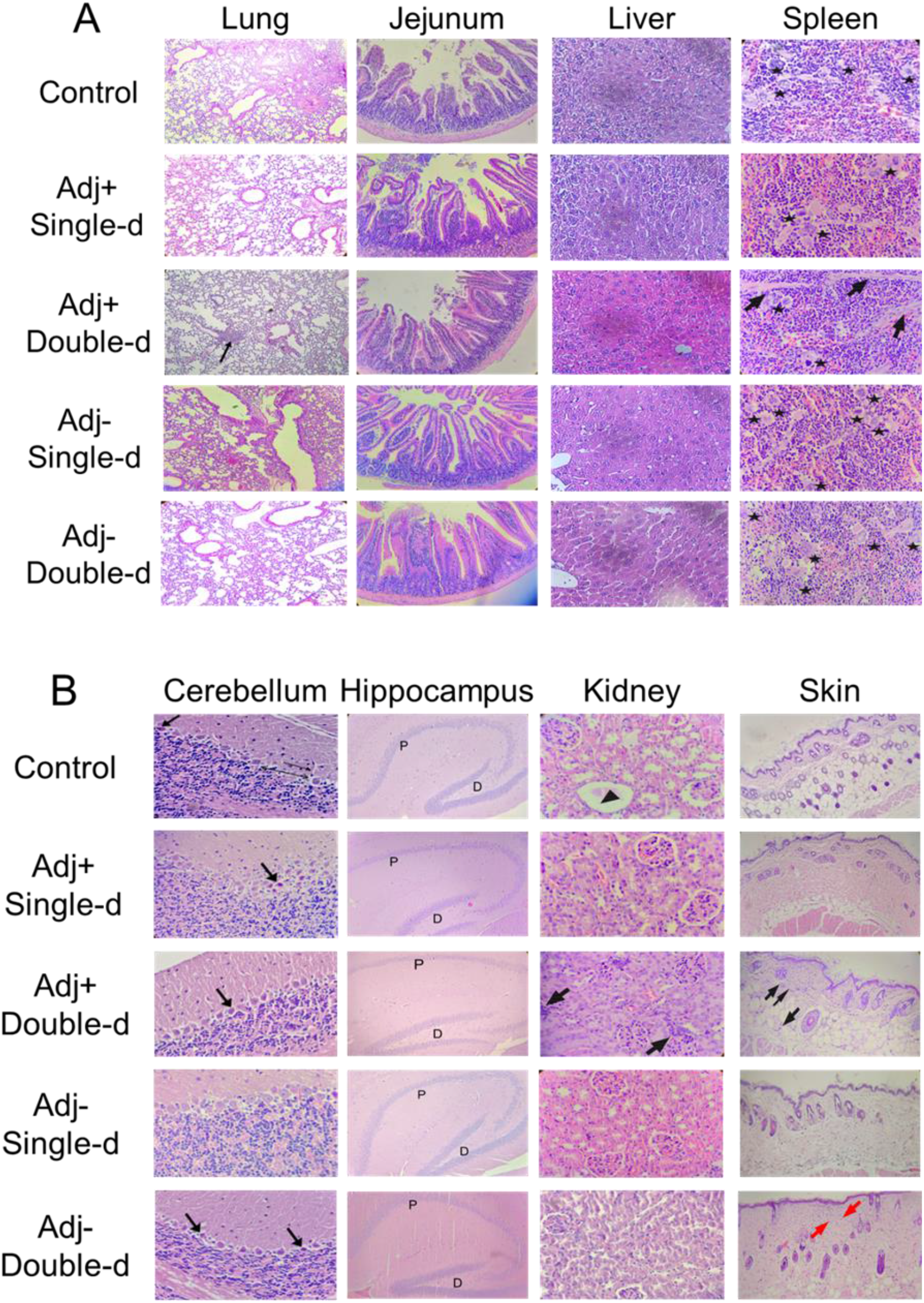
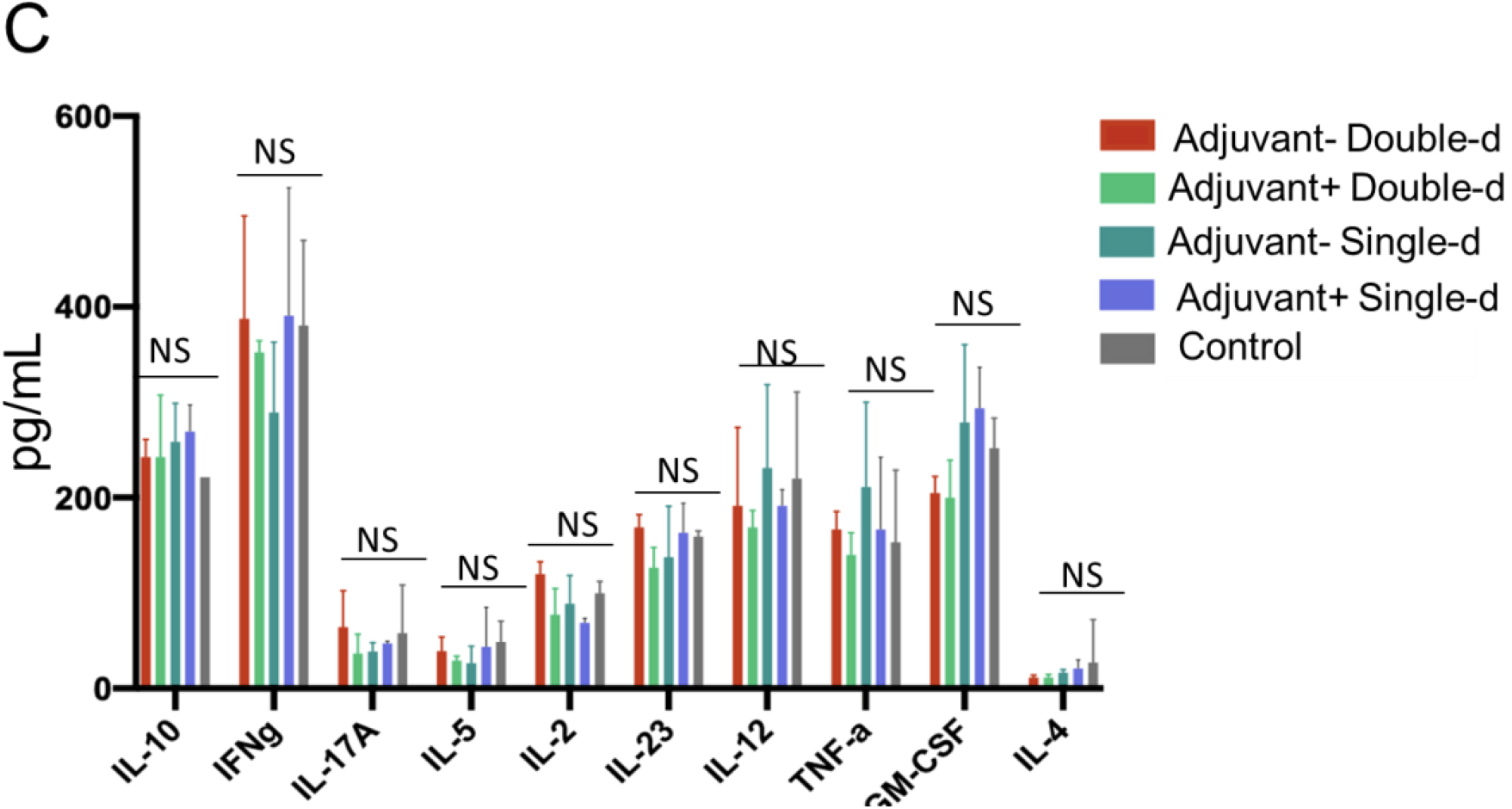
Day-21 Safety analysis of the vaccine candidates, SK-01 V1, and OZG-3861 V1. Histopathologic analysis on day 21 of **A.** lung (the arrow; chronic inflammation X100), jejunum of the intestine, liver, spleen, and **B.** cerebellum of brain, hippocampus, kidney, and skin. In the spleen, stars show foci of megakaryocytes and arrows were trabeculae (X400 H+E stain). In the cerebellum, the thin arrow shows dendrites, the thick arrow shows picnotic cell, the thin double-headed arrow shows space. In the hippocampus, P is the pyramidal cell layer and D is the dentate gyrus (X100 H+E stain). In the kidney, the arrow shows interstitial chronic inflammation (X400 H+E stain). In the skin, black arrows show infiltrated inflammatory cells and red arrows show eosinophils (X40 H+E stain). **C.** Bar graphs showing quantitated mouse cytokine bead array analysis by assessing Th1, Th2, and Th17 specific cytokines (picogram/ml) in mice serum collected on day 21 of the vaccine treatment.

### Pre-clinical efficacy and dose study of vaccine candidates, SK-01 V1 and OZG-3861 V1

In order to perform in vivo efficacy and dose studies of vaccine candidates, OZG-3861 V1 and SK-01 V1 were administered intradermally to the mouse groups as single or 15-day booster doses (**Fig. 5A**). SARS-CoV-2 specific IgG antibody analysis was performed in three different titrations (1:64, 1: 128, and 1: 256) in serum isolated from blood. According to the IgG ELISA result, a significantly increased SARS-CoV-2 IgG antibody was detected in all groups in comparison with the control (non-vaccinated) mouse group (*p<0.05) **(Fig. 7A).** Mouse SARS-CoV-2 Spike S1 monoclonal antibody was used in the same test as a positive control for the accuracy of the analysis **(Fig. 7A).** The proof-of-concept has been optimized with the real-time cell analyzing (RTCA) system to determine the neutralization efficiency of SARS-CoV-2 specific antibodies in serum content. As a representative data, with double dose SK-01 V1, control group serums were pre-incubated with SARS-CoV-2 virus at 100x TCID50 dose **(Fig. 7B)** in 1: 128 and 1: 256 ratios, followed by Vero cell viability for approximately 6 days according to normalized cell index value in the RTCA system **(Fig. 7C).** The results showed that double dose SK-01 V1 can neutralize the infective virus significantly in comparison with the control serum group even at 1: 256 dilutions **(Fig. 7C).** Conduction of the same study for a single dose of SK-01 V1 and a single and double dose of OZG-3861 V1 showed that double dose OZG-3861 V1 at 1: 256 dilution also had virus neutralization capacity **(Fig. 7D).** Although single-dose SK-01 V1 or OZG-3861 V1 did not show a significant neutralization efficacy at 1: 256 dilution (p>0.05), it was evaluated that it had neutralization capacity at 1: 128 dilution **(Fig. 7D).** The high rate of neutralizing antibodies detected in control mice (in 2 out of 3 mice, %66) in this study suggests that the mice may have previously had a viral infection like the mouse hepatitis virus (MHV), a member of the coronavirus family, related to SARS-CoV-2. Therefore, the findings of this study show the need to repeat the assay with mice that were considered to be negative for spontaneous neutralizing antibodies. However, the ADE (antibody-dependent enhancement) test worked almost like a confirmation of the neutralizing antibody test, showing that the antibodies formed neutralized the virus without causing ADE **(Fig. 7E).** This in vitro analysis with mice serum showed that the SARS-CoV-2 specific neutralizing antibody was produced with the help of immunization of mice with the first versions of SK-01 and OZG-3861 vaccine candidates without an ADE effect.

**Figure 7.**
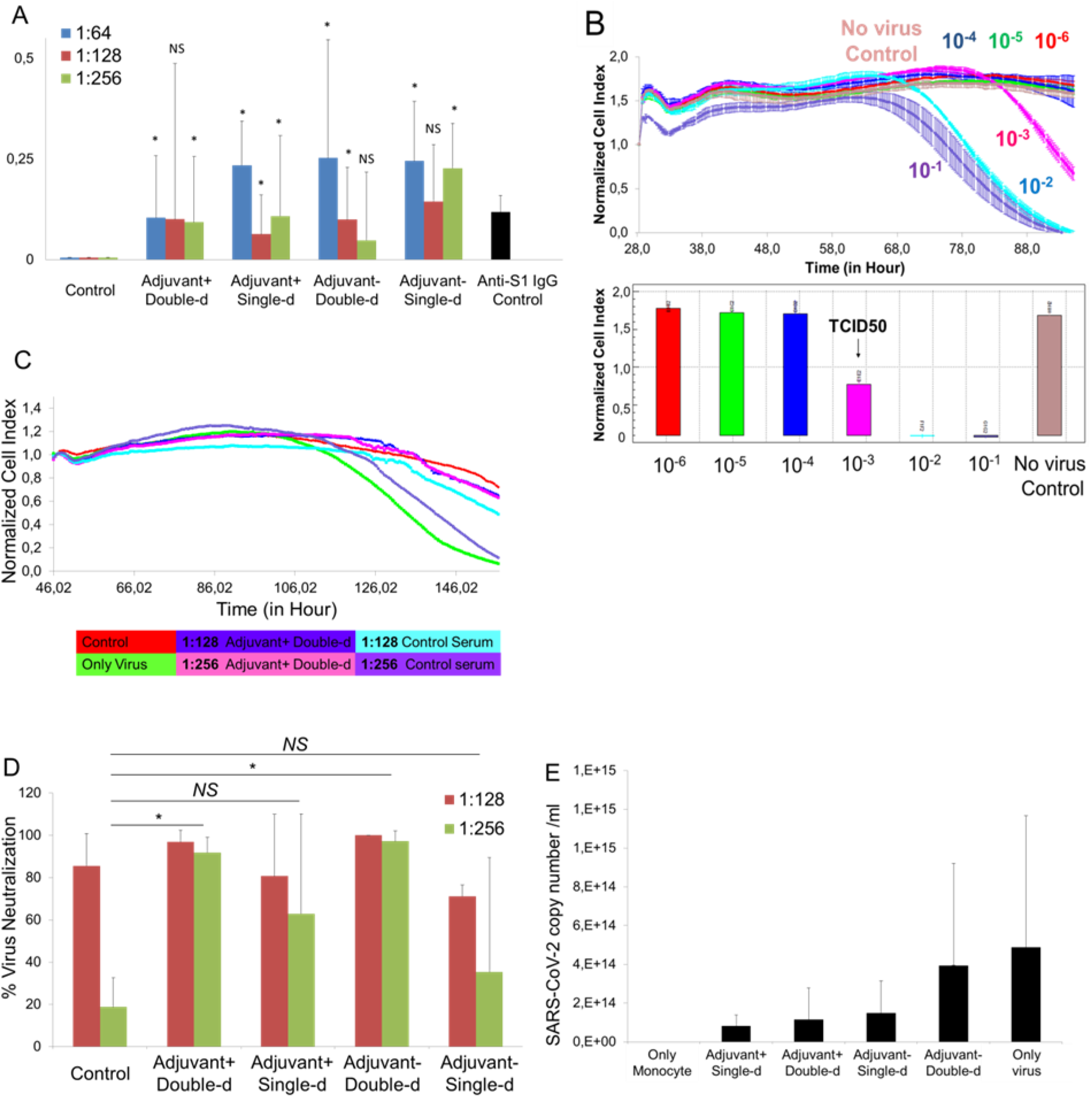
Pre-clinical efficacy study of vaccine candidates, SK-01 V1, and OZG-3861 V1. **A.** Bar graph showing SARS-CoV-2 specific IgG analysis of the groups concerning the positive control antibody, mouse SARS-CoV-2 Spike S1 monoclonal antibody using ELISA. **B.** Upper graph showing RTCA analysis of infective active SARS-CoV-2 in a dose-dependent manner for 6 days. Bar graph showing quantified normalized cell index values of SARS-CoV-2 titrations to determine TCID50 dose. **C.** Representative RTCA graph of neutralization assay in which 1:128 and 1:256 dilutions of adjuvant positive double-dose (SK-01 V1) and control mice serum preincubated with 100x TCID50 dose of SARS-CoV-2. **D.** Bar graph showing quantified virus neutralization ratio of the vaccine treated groups at 1:128 and 1:256 dilutions. **E.** Bar graph showing quantitated SARS-CoV-2 copy numbers when culturing on healthy adult monocytes along with 1:256 mice serum to determine Antibody-Dependent Enhancement (ADE). The threshold of significance for all tests was set at *p<0.05. *NS* is Non-Significant.

In this study, following the finding of the presence of antibodies due to B cell activity, T cell response was tested upon re-stimulation either with whole inactivated SARS-CoV-2 virus or SARS-CoV-2 specific S, N, and M-protein peptide pool. T cells were isolated from the spleen tissue of mice dissected either on day 21 or day 34. As T cells were incubated with the SARS-CoV-2 antigens, the cytokine secretion profile was evaluated for 72 hours (**Fig. 8A**). Subsequently, the balance of Th1 and Th2 cell responses was determined and showed an increase in the ratio of IL-12 to IL-4 and IFNγ to IL-4 (**Fig. 8B**). This illustrated that our vaccine candidates were predominantly biased towards Th1 CD4 T cell response regarding control T cells isolated from untreated mice spleens. Furthermore, a significant increase in the proportion of cytotoxic CD8 T cells from an adjuvant negative single dose (OZG-3861 V1) and adjuvant positive double dose (SK-01 V1) immunized mice upon re-stimulation with the peptide pool was detected (**Fig. 8C**). This showed that viral antigens caused CD8 T cell proliferation 34 days after vaccination. However, there was no increase in the T cell activation marker, CD25 on both T cell subtypes (**Fig. 8C**). In order to evaluate the SARS-CoV-2 specific T cell response, stimulated T cells providing specific IFNg secretion that were counted as spots in the IFNγ ELISPOT plate were analyzed (**Fig. 8D and 8E**). Findings especially showed that IFNg increase in T cells isolated from day21-dissected spleens was in the single or double dose of SK-01 V1 vaccine candidate containing adjuvant as opposed to the control mouse group (**Fig. 8D**). Although there was no significant difference in the double dose of the OZG-3861 V1 vaccine candidate without adjuvant, an increase in IFNg was detected in single-dose administration (**Fig. 8D**). On the other hand, a significant IFNg secretion from the T cells isolated from day34-dissected spleens of the single or double dose of OZG-3861 V1 and a single dose of SK-01 V1 was detected (**Fig. 8E**). This analysis illustrated that SK-01 V1 and OZG-3861 V1 vaccine candidates can achieve not only B cell response but also T cell response. These encouraging pre-clinical in vivo efficacy studies will lead us to challenge humanized ACE2 + mice with SK-01 V1 and OZG-3861 V1.

**Figure 8.**
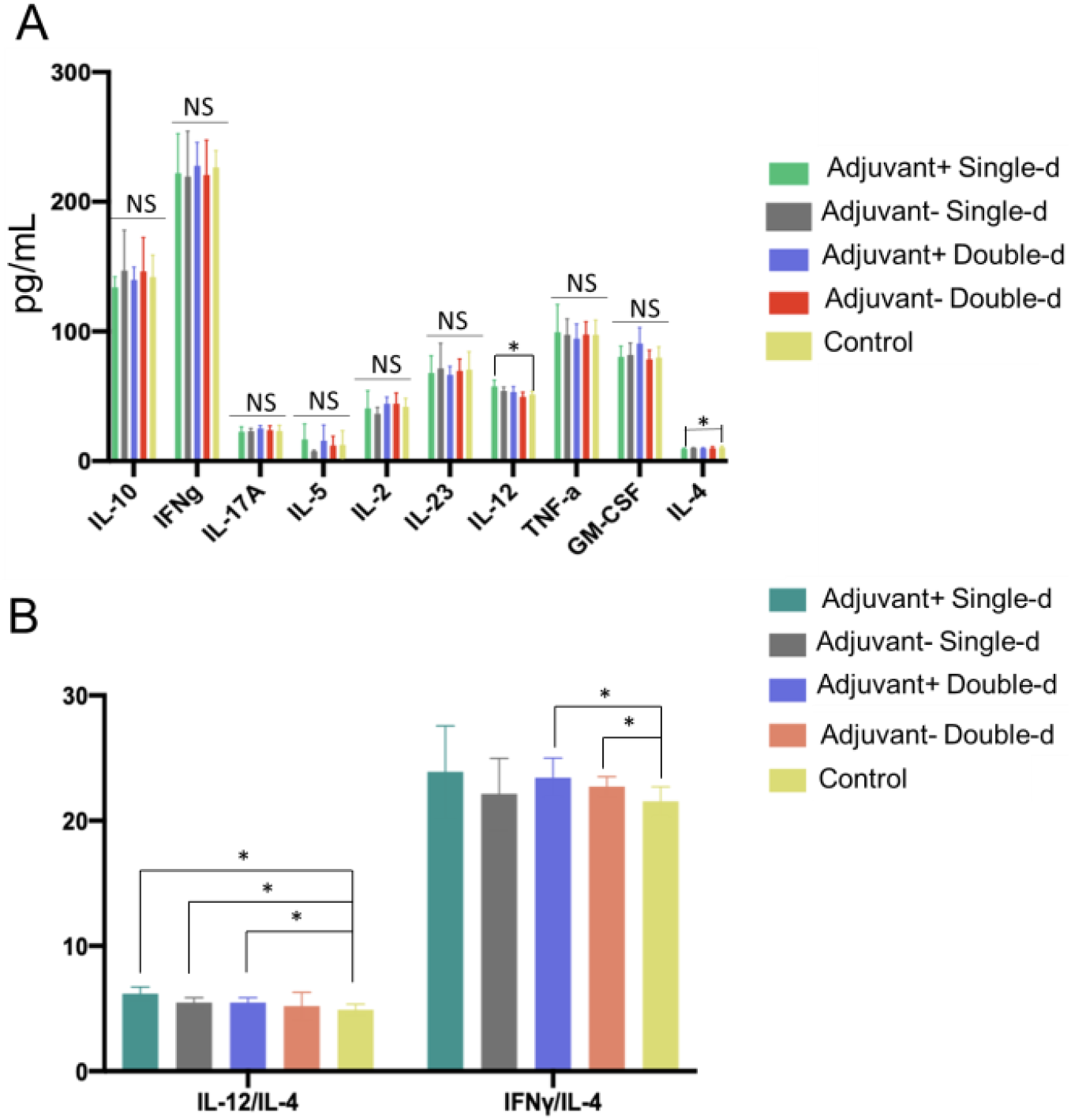
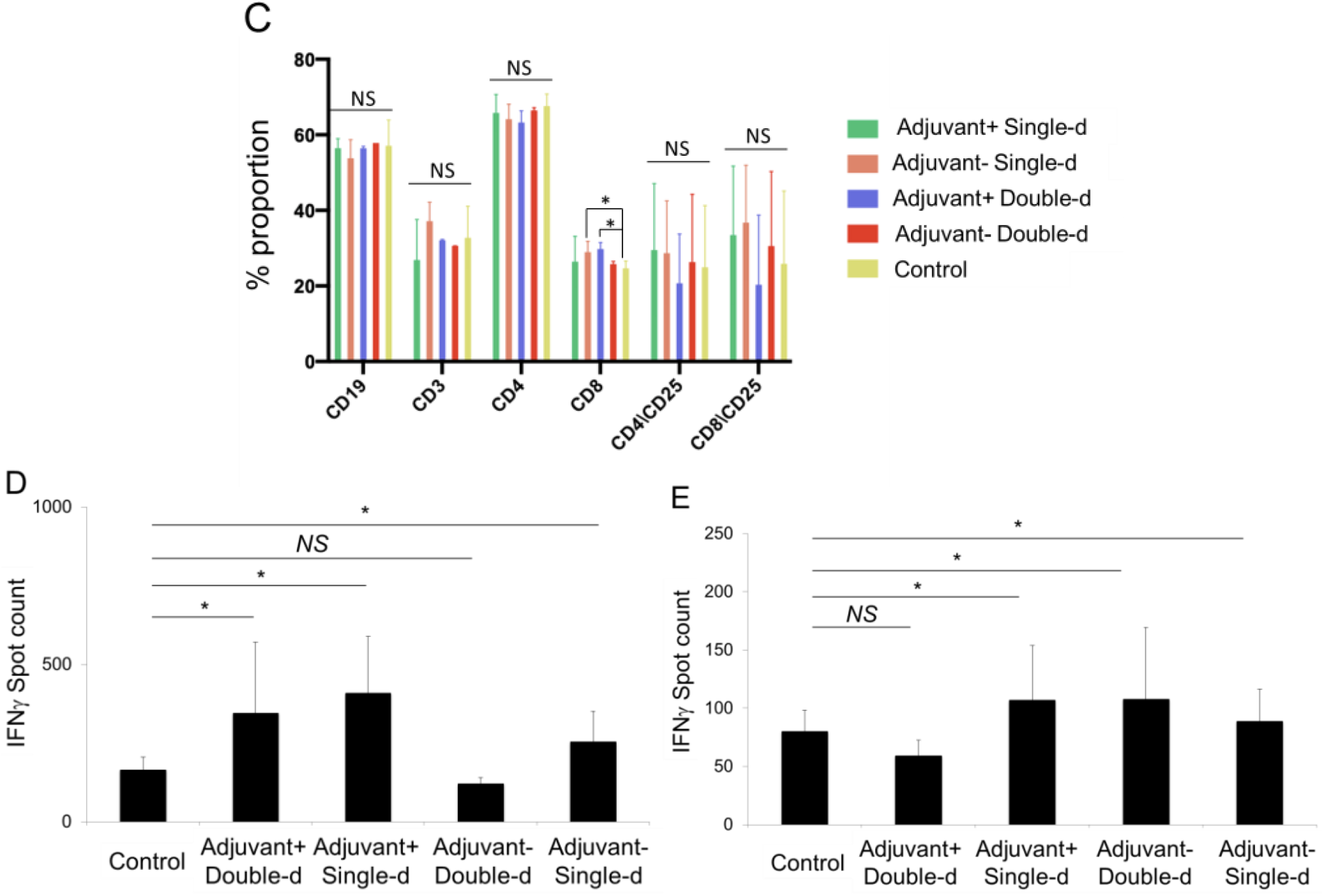
Mouse Spleen T cell response upon SARS-CoV-2 antigen. **A.** Bar graphs showing quantitated mouse Th1, Th2, and Th17 specific cytokines (picogram/ml) secreted by the T cells isolated from dissected spleens on day 34 re-stimulated with SARS-CoV-2 specific S, N, and M-protein peptide pool. **B.**Bar graphs showing the balance of Th1 and Th2 CD4 T cell response. The ratio of IL-12 to IL-4 or IFNg to IL-4 demonstrates Th1–dominant response. **C.** Bar graph showing a change in the proportion of immune cell subtypes re-stimulated with the peptide pool (B cell, CD19+; T cell, CD3+; T helper cell, CD3+ CD4+ and cytotoxic T cell, CD3+ CD8+). The activation marker of T cells is the upregulation of CD25. **D.** Bar graph showing IFNγ spots formed during mouse spleen T cells isolated on day 21 incubated with whole inactive SARS-CoV-2 virus for 72 hr. **E.** Bar graph showing IFNγ spots formed during mouse spleen T cells isolated on day 34 incubated with SARS-CoV-2 specific S, N, and M-protein peptide pool for 72 hr.

## Discussion

Various methods are available in obtaining an effective and safe immunization in inactive vaccine production. Besides chemical modifications such as formaldehyde or β-propiolactone, physical manipulations with ultraviolet radiation or gamma radiation are also available ^9^. Chemical modifications in the vaccine inactivation process are time-consuming methods due to the need for purification. They also have disadvantages associated with toxicity with changes in viral structure and product loss due to the necessity to purify the final product. The physical inactivation process in the three separate animal experiments in this study showed that single or double dose administrations with or without adjuvant were non-toxic and effective.

All viral vaccines contain virus-like materials that they try to protect. This directs the immune system to generate a response and to produce antibodies ready for use if it encounters a true viral infection. However, it is worrying that the virus mutates to form "escape mutants". These are mutated versions of the virus that vaccine-induced antibodies do not recognize. For a significant immunization, it is necessary to create a vaccine profile that covers the genetic variation of the virus within the community. If the genetic variation represented by the vaccine is small, triggering social immunity will not be at the desired rate. Therefore, the production and selection of more than one inactive viral strain remain an important mechanism for producing successful viral vaccines. Different variations may occur when producing large quantities (bulk) in the laboratory. Due to the low sensitivity of RNA-bound RNA polymerase, RNA viruses always produce a pool of variants during replication ^10^. This phenomenon provides a potential for the rapid evolution of the virus, but it also makes up the majority of mutations that have detrimental effects on virus stability. Increased virus complexity can cause a weakening of the population's virulence degree; therefore, the characterization of individual variants can provide useful information for the design of a new generation of more effective and safer vaccines. Lyophilization (freeze-drying) is expressed as a process that combines the benefits of freezing and drying to obtain a dry, active, long shelf life, and easily soluble product^11^, and it is an important process for the preservation of heat-sensitive biological materials ^12^.

In the production of inactive vaccines in this study, both inactivation and sterilization steps were carried out with a double dose (fractionated) gamma irradiation (25 kGy / single dose). With gamma irradiation, the frozen product can be irradiated, thus reducing the risk of toxicity as a free radical formation is prevented, and the risk of possible changes in the viral protein structure is reduced. Since functional viral structures are preserved, both B cell and T cell responses are triggered. With the first dose of irradiation, the raw product containing the live and infective virus is transformed into an intermediate product containing an inactive virus. Thus, prior to bottling, both environment and personnel are protected. A radiation dose over 25 kGy single dose was sufficient for inactivation is reported ^13^. Both RTCA assay and colorimetric MTT assay for cell viability and proliferation confirmed the inactivation of the virus propagated in this study following 21-day of incubation with the three independent inactivated virus samples, and no suspicious situation was observed. In this study protocol, following the conversion of the frozen raw product into an inactive form by gamma irradiation, it is melted, bottled, and lyophilized. Lyophilized formulations, together with the advantage of better stability, provide easy handling during transportation and storage. The second dose of irradiation, performed after bottling (vialing) and lyophilization, functions to eliminate the presence of the replicant virus and end product sterilization. Also, unlike chemical inactivation methods, isolation and purification processes are not required. As a result, while achieving inactivation and sterility, fractionated (2-stage) gamma irradiation leads to less damage to the final product virus structure and allows the maximization of the preserved products. Besides, in our recently published pre-print ^14^, we optimized an inactivated virus vaccine that includes the gamma irradiation process for the inactivation as an alternative to classical chemical inactivation methods so that there is no extra purification required. Also, we applied the third version of our vaccine candidate (OZG-38.61.3) using the intradermal route in mice which decreased the requirement of a higher concentration of inactivated virus for proper immunization unlike most of the classical inactivated vaccine treatments ^14–16^. In this study, we immunized human ACE2-encoding transgenic mice and infected them with a dose of infective SARS-CoV-2 virus for the challenge test. We showed that the vaccinated mice showed lowered SARS-CoV-2 viral copy number in oropharyngeal specimens along with humoral and cellular immune responses against the SARS-CoV-2, including the neutralizing antibodies ^14^.

In toxicity analysis of vaccinated mice in this study, it was decided that adjuvant positive double dose administration should be removed in the newly designed version 2 vaccine model due to the finding of inflammatory reaction in the skin and kidney. Furthermore, for the overall picture immunization in the presence or absence of GM-CSF adjuvant did not yield significant differences in antibody and T cell responses. With this study, no significant difference was observed on immunization when GM-CSF was used as an adjuvant. Studies also showed that injection with intradermal GM-CSF leads to significant increases in grafting power in intradermally applied areas compared to distant areas ^17^. This may explain the inability of GM-CSF in intradermally administered inactive virus vaccines. Therefore, it was decided to increase the SARS-CoV-2 effective viral copy dose (1×10^13^ or 1×10^14^ viral copies per dose) in version 2 of vaccine candidates. In terms of T and B cell responses, it was observed that especially the vaccine models containing GM-CSF as an adjuvant lead to significant antibody production with neutralization capacity in the absence of ADE feature.

On the other hand, ACE2 is ubiquitously expressed in various types of cells in humans ^18^. Interaction between SARS-CoV-2 and ACE2 stimulates various pathways which some of which are known to determine the pathogenesis of SARS-CoV2 infection ^19^. It was reported that ACE2-mediated cardiovascular protection was lost, multiorgan failure and gut dysbiosis were taken place following endocytosis of the enzyme following interaction with SARS-CoV-2 ^20^. However, we applied the vaccine candidate (OZG-38.61.1) using the intradermal route in mice which decreased the requirement of a higher concentration of inactivated virus for proper immunization unlike most of the classical inactivated vaccine treatments ^15, 16^. Therefore, we expected that only specialized dendritic cells named Langerhans cells (LCs) that populate the epidermal layer of the skin would be primed with the inactivated SARS-CoV-2 particles in the site of injection for the proper immunization ^21^. Furthermore, in the histopathological analysis that was performed in this study and our recent preprint ^14^, we did not determine any signature of multiorgan failure or cardiovascular impairment, supporting the safety of our vaccination procedure.

In the formation of the SARS-CoV-2 specific antibody, the antibody was detected up to 1: 256 titration in all doses and formulations of vaccine candidates administered to mice. On the other hand, in the control mice, we determined a low level of spontaneous neutralizing antibodies, which may be because the control mice may meet coronavirus like infections such as mouse hepatitis virus (MHV) ^22, 23^. Also, no traces of MHV was detected when stool samples from 5 mice were tested for MHV copy using qRT-PCR. This neutralizing antibody ratio was seen in the first version of vaccine candidates with a viral copy number of approximately 10^7^/dose is predicted to achieve higher and longer-term antibody concentration in the second version which will have 9×10^11^ or 1×10^13^ viral copies per dose of SK-01 V2 and OZG-3861 V2 vaccine candidates. To assess the sensitivity or specificity of the XCELLIGENCE assay, we tested the OZG-38.61.3 vaccine candidate in our recent challenge study with ACE2 transgenic mice. No significant neutralizing capacity was observed in the neutralizing test using SARS-CoV2 at the same proportions with convalescent plasma or standard control sera ^14^.

We also determined the balance of Th1 and Th2 cell responses, because Vaccine-Associated Enhanced Respiratory Disease (VAERD) was reported to be associated with Th2-biased cell responses, both in some animal models and children vaccinated with whole-inactivated RSV and measles virus vaccines ^24–27^. In this study, Th1 to Th2 was balanced towards Th1 response suggesting that VAERD risk is low. Another finding showed that the presence of adjuvant is more important in T cell response in comparison with the B cell response, which may lead to long-term immunization. In addition, here, the ADE test was the in vitro equivalent of VAERD. It has been reported indirectly that there is no vaccine-related ADE effect within the macrophage. The absence of this effect has been confirmed in Version 3 of our vaccine OZG-38.61 in challenge tests with ACE2 transgenic mice ^14^. VAERD evolves the ADE in in-vitro conditions. It has been reported in our latest report indirectly that there is no vaccine-related ADE effect within the macrophage. The absence of this effect has been confirmed in the Version 3 challenge tests ^14^. In the viral challenge study, viral dissemination was blocked by SARS-CoV-2 specific antibodies and neutralizing antibodies in both vaccine groups, unlike other groups, especially at high doses, CD4 activation is also present in the immune response that occurs. the resulting T cell response is in the Th1 response type as desired; the resulting T cell response is in the Th1 response type as desired. It has been determined that the ADE effect is not observed. These findings also confirm that our vaccine is non-replicative ^14^.

With this study, it was seen that our gamma-irradiated inactivated vaccine candidates can effectively trigger the production of SARS-CoV-2 specific antibodies along with long-term T cell response. Hence, the findings of this study prompted us to plan a new in vivo experiment with the second version of SK-01 and OZG-3861 (1×10^13^ or 1×10^14^ viral copies per dose) in humanized ACE2+ mice ^14^. In this report, we determined that GMCSF adjuvant positive vaccine administration should be removed in the newly designed version of the OZG-38.61 vaccine model due to the finding of inflammatory reaction in the skin, cerebellum, and kidney in toxicity analysis of vaccinated mice. Therefore, it was decided to increase the SARS-CoV-2 effective viral copy dose (1×10^13^ or 1×10^14^ viral copies per dose) in the last version of vaccine candidates. In the challenge study, we produced the third and final version of the OZG-38.61 without an adjuvant ^14^. In this study, it was demonstrated that the OZG-38.61.3 vaccine candidates that we created with gamma-irradiated inactivated SARS-CoV-2 viruses produced neutralizing antibodies, especially effective in 10^14^ viral copy formulation, and this was effective in transgenic human ACE2 expressing mice. We showed that it can protect against infection ^14^. This preclinical study has encouraged us to try phase 1 vaccine clinical trials to avoid the COVID-19 pandemic.

## Acknowledgment

Sevda Demir and Sezer Akyoney are funded by TÜBİTAK 2247-C Trainee Researcher Scholarship Program (STAR). Selen Abanuz and Ilayda Sahin are supported by the TUBITAK-BIDEB 2244-University-Industry Ph.D. program, project number 118C082. Figure 1 and 5A of the study was drawn by G.S.K. and C.T. with BIORENDER.com.

