## Supplemental Figure 1 for "Preclinical efficacy and safety analysis of gamma-irradiated inactivated SARS-CoV-2 vaccine candidates"

supplementary figure

### Supplementary Figure 1

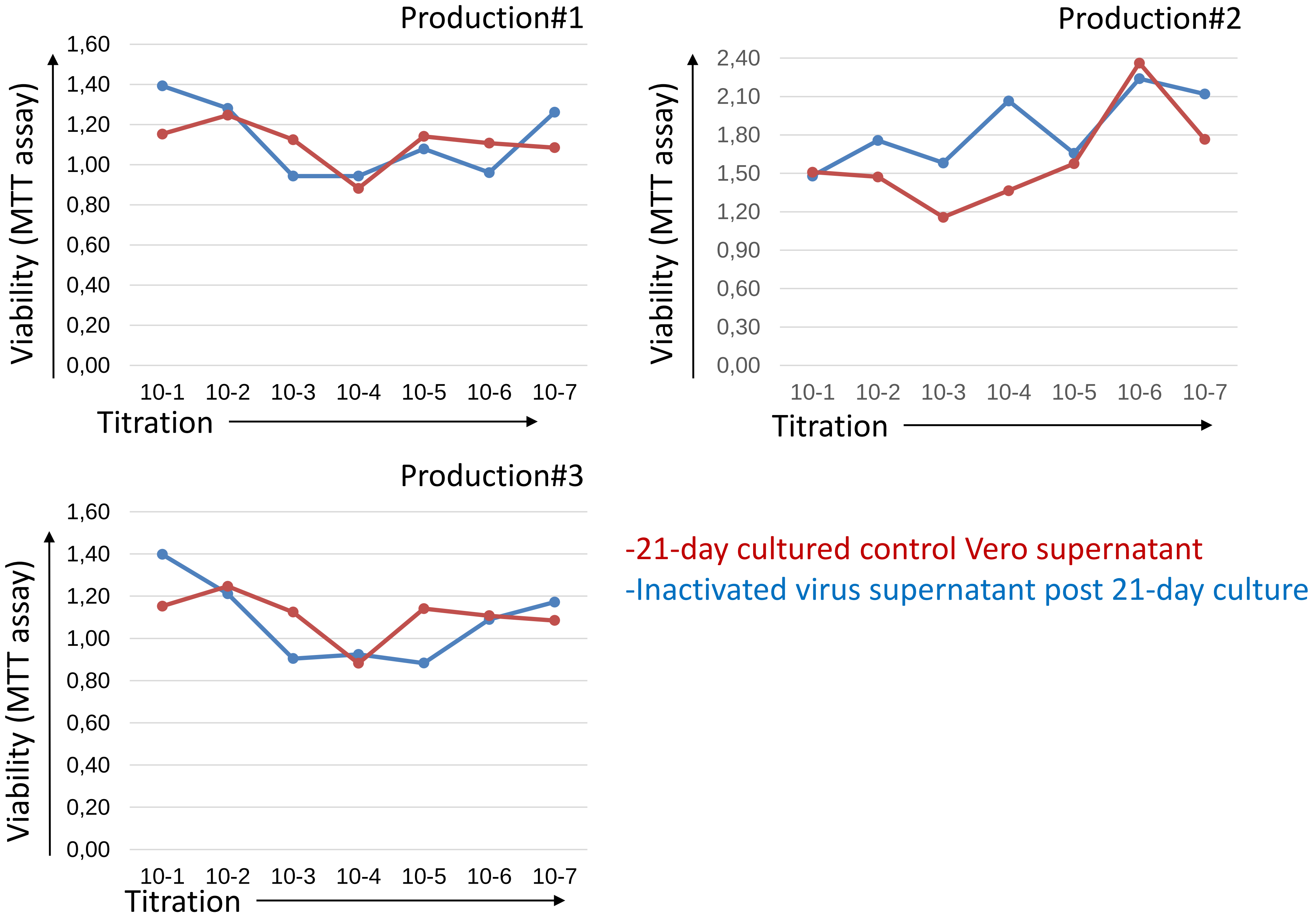

**Supplementary Figure 1:** Colorimetric MTT viral culture assay. Three independent inactivated virus production was tested for the inactivation by inoculating Vero cells for 21-day of culturing with subsequent passages using 1:2 refreshing culturing media. At the end of the 21-day culturing, the active virus was expected to be propagated if existed in the vaccine samples. To assess the cytopathic effect of the potential infective virus, an MTT assay was performed as described 14. The Redline is to control Vero supernatant post 21-day culturing. The blue line is the supernatant of Vero cells inoculated with inactivated Virus vaccine sample post 21-day culturing.
