## Supplemental Table 1 for "Preclinical efficacy and safety analysis of gamma-irradiated inactivated SARS-CoV-2 vaccine candidates"

**Supplementary Table 1: Quality Control Limits and Tests Definition and Result Table For COVID-19 Vaccine**

| **VERO CELL BEFORE COVID-19 VACCINE VIRUS CULTURE** | | | | | | |
| --- | --- | --- | --- | --- | --- | --- |
| **PARAMETERS** | **METHODS** | **LIMITS** | **EXPLANATIONS** | **RESULTS** | **REFERENCES/CRITERIA** | |
| Number of Passages | Registration / Label | 16-100 Passages | Documents and labels showing ATCC or place/date of receipt. | ATCC CCL-81 VERO Cell  Lot Number: 70016956  Number of Passages: 13th Passage | (Barrett et al., 2009; Hoeksema et al., 2018; Hu et al., 2015; Kistner et al., 1998; Shen et al., 2019) | |
| Viability | Trypan Blue | > %90 | - | Viability of 13th passage: %97 | (Felice et al., 2009) | |
| Intact Live Cell Microscopy and Cell Culture | Microscope Culture | > 90% Confluency | The cells should appear intact and maintain integrity.  5x10^6^ Vero cells were cultured in a T25 flask and 90% cell occupancy rate at the end of 24 hours (confluency). | Integrity observed.  > 90% Confluency | (Ammerman et al., 2008) | |
| Cytopathic Effect | Real-Time Cell Analysis (RTCA) / MTT Based Cytopathic Effect Analysis | Spontaneous: Similar Properties  Healthy Cell: Should Not Show Cytopathic Effect | Should show similar culture characteristics on day 7 with HEK and/or MSC cells. (Spontaneous)  VERO lysate should not show cytopathic effect in HEK / MSC / Vero cells on the 7th day. (Healthy) | It showed similar properties in the spontaneous cytopathic effect.  No cytopathic effect was observed in healthy cytopathic effect. | EP-01/2008:0443  WHO/BS/10.2132  (Page:60) | |
| Chromosome Analysis | Cytogenetics | 54-61 Chromosome | - | 57-59 chromosomes observed. | EP-01/2017:50203 | |
| Viral Serology (HBV, HCV, HIV…) | PCR / Cytopathic Effect Analysis / TEM Analysis | NEGATIVE | Adeno Virus, Herpes Virus, HIV, Hepatit B Virus, Hepatit C Virus, BK Virus, Influenza A/B Virüs, Bovine Respiratory Syncytial Virus,  Infectious Bovine Rhinotracheitis Virus, Parainfluenza Virus, Rabies Virus.  Also, TEM images and healthy cells are confirmed by a cytopathic effect. | Adenovirüs DNA PCR: Negative  EBV DNA PCR: Negative  HIV RNA PCR: Negative  HBV DNA PCR: Negative  HCV RNA PCR: Negative  BK Virüs DNA PCR: Negative  Influenza A/B RNA PCR: Negative  HIV Antikoru 1&2 + p24 Antijeni: Negative  BRSV: The result is expected.  BVDV: The result is expected.  IBR: The result is expected.  PI3: The result is expected.  RABIES: The result is expected. | EP-01/2017:50203  WHO/BS/10.2132  (Page:60) | |
| Purity | TEM | NEGATIVE | Pathogen Virus Analysis | Negative | EP-01/2017:50203  WHO/BS/10.2132  (Page:60) | |
| Microbiological Quality Control | Culture | NEGATIVE | - | Negative | 2.6.1 (method of reference)  EP-01/2017:50203 | |
| Fungi Culture | Culture | NEGATIVE | - | Negative | 2.6.1 (method of reference)  EP-01/2017:50203 | |
| Mycoplasma | PCR | NEGATIVE | - | Negative | 2.6.7 (method of reference)  EP-01/2017:50203 | |
| **COVID-19 VACCINE VIRUS CULTURE INTERIM PRODUCT (VIRAL INTERIM STOCK)** | | | | | | |
| **PARAMETERS** | **METHODS** | **LIMITS** | **EXPLANATIONS** | **RESULTS** | **REFERENCES** | |
| Amount of Virus | RT-qPCR | > 1x10^11^ | The method is done to see the amount of virus produced. | Desired dose provided > 1x10^15^ | Producer references are sufficient. | |
| Microbiological Quality Control | Culture | NEGATIVE | - | Negative | 2.6.1 (method of reference)  EP-01/2017:50203 | |
| Fungi Culture | Culture | NEGATIVE | - | Negative | 2.6.1 (method of reference)  EP-01/2017:50203 | |
| Mycoplasma | PCR | NEGATIVE | - | Negative | 2.6.7 (method of reference)  EP-01/2017:50203 | |
| **COVID-19 VACCINE (VIRAL MAIN STOCK)** | | | | | | |
| **PARAMETERS** | **METHODS** | **LIMITS** | **EXPLANATIONS** | **RESULTS** | **REFERENCES** | |
| Number of Passages | Registration / Label | < 15th Passage | Documents and labels showing ATCC or place/date of receipt. | 6-9 Passages | (Barrett et al., 2009; Hoeksema et al., 2018; Hu et al., 2015; Kistner et al., 1998; Shen et al., 2019) | |
| Amount of Virus | RT-qPCR | > 1x10^15^ /ml (RT-qPCR) | - | 3,5 x 10^20^ x 300ml | (Krammer, 2020) | |
| Amount of Virus | Real-Time Cell Analysis (RTCA) / MTT Based Cytopathic Effect Analysis | > 1x 10^6^ TCID50 | - | 10^14^ için > 4 TCID50/dose  10^15^ için > 40 TCID50/dose | (Stefanowicz-Hajduk & Ochocka, 2020) | |
| Genotype Analysis | Sequencing | A seed must be >98% compatible with the genetic sequence | It should also include> 95% of existing mutations.  It is recommended to do 1 in 10 passages. | In the viral gene pool of 10,000 nucleotides, 15 nucleotide changes were observed after the 9th passage. This change rate is 0.1%.  It includes all defined mutations. | EP-01/2010:51400  (Page:871)  (Bar-On et al., 2020) | |
| Identity | RT-PCR | POSITIVE | - | Positive | EP-04/2009:2308 | |
|  | TEM | POSITIVE | Oval particles include spikes that size is 70-160 nm. | Positive | (Prasad et al., 2020) | |
| Purity | VERO Protein ELISA | < 5 mg/ml | - | 0,35 mg/ml | The residual Vero cell host protein content of 6 batches of JE vaccine prepared with Vero cells was 153.3-5 850.9 ng/ml, and that of 5 batches of rabies vaccine prepared with Vero cells was 1 895.7-40 625.1 ng/ml.  (Jia et al., 2009) | |
|  | VERO DNA NanoDrop | < 1mg/ml | - | 0,158 mg/ml | EP-04/2009:2308  WHO established the acceptable limit of not more than 100 μg of cellular DNA per human dose.  WHO Expert Committee on Biological Standardization Fifty-sixth Report (A.4.3.4.4) | |
| Biochemical Analysis | pH, K, Ca, Glucose, Total Protein, Albumin | pH: 6-8  Glucose:<1mg/ml  K:< 1mg/L  Ca:< 1mg/L  Total protein: < 1g/dl  Albumin: < 0.6g/dl | - | pH:7  K: 2,01 mmol/L  Ca: 1,40 mg/dL  Glucose:24 mg/dl  Total Protein:1 g/dl  Albümin:0.6 g/dl | **Potentiometric determination of pH (2.2.3 method reference):**  EP-07/2016:20203  **Cl (2.4.4 Chloride):**  EP-01/2008:20404  **K (2.4.12 Potassium):**  EP-01 /2008:20412  **Ca (2.4.3 Calcium):**  EP-01/2008:20403  **Glucose:**TFA-01/2017:20270 (Page:473)  The albumin content which ideally should not exceed 1% of total protein content.  Total Protein-Albumin: WHO/BS/2016.2300 – (Page:89 – 34th Line) | |
| Microbiological Quality Control | Culture | NEGATIVE | - | Negative | 2.6.1 (method reference)  EP-01/2017:50203 | |
| Fungi Culture | Culture | NEGATIVE | - | Negative | 2.6.1 (method reference)  EP-01/2017:50203 | |
| Mycoplasma | PCR | NEGATIVE | - | Negative | 2.6.7 (method reference)  EP-01/2017:50203 | |
| Endotoxin | LAL Test | <5 U/dose | - | Negative | EP-01/2010:50110 (Page:712) | |
| Stability / Effectiveness | Cytopathic Effect with TCID50 | POSITIVE | Efficiency analysis of 0,1,6,12,24 and 26th months under <-60 ˚C storage conditions. | 1th Month Positive / No Change | EP-01/2017:0062 (Page:961) | |
| **COVID-19 VACCINE VIRUS INTERIM PRODUCT AFTER INACTIVATION (INACTIVE MAIN STOCK)** | | | | | | |
| **PARAMETERS** | **METHODS** | **LIMITS** | **EXPLANATIONS** | **RESULTS** | **REFERENCES** | |
| Identity | Single Radial Immunodiffusion (SRID) | POSITIVE | Identity with SARS-CoV-2 specific S protein antibody. | Positive | 2.7.1 (method reference)  EP-01/2008:0158  EP-04/2009:2308 | |
|  | TEM | POSITIVE | Oval particles include spikes that size is 70-160 nm. | Positive | (Prasad et al., 2020) | |
| Size and Particle Analysis | NanoSight | Size: 60-200 nm  Particle: > 1x10^8^ ml | - | Size: 228.3 +/- 5.3 nm  (200 nm above indicates the presence of aggregate.)  Particle: 1,26e+10 ± 1,10e+09 particles/ml | Size Analysis:  (Huang et al., 2020)  Particle Analysis:  (Zhu et al., 2020) | |
| Biochemical Analysis | pH, Na, Cl, K, Ca, Glucose, Total Protein, Albümin | pH:6-8  Glucose: <1mg/ml  K: <1mg/L  Ca: <1mg/L  Total protein: <1g/dl  Albumin: < 0.6g/dl | - | pH:7  K: 0,08 mmol/L  Ca: 0,1 mg/dL  Glucose:1 mg/dl  Total Protein:<0,1 g/dl  Albümin:<0.6 g/dl | Potentiometric determination of pH (2.2.3 method reference):  EP-07/2016:20203  Cl (2.4.4 Chloride):  EP-01/2008:20404  K (2.4.12 Potassium):  EP-01 /2008:20412  Ca (2.4.3 Calcium):  EP-01/2008:20403  Glucose:  TFA-01/2017:20270 (Page:473)  The albumin content which ideally should not exceed 1% of total protein content.  Total Protein-Albumin: WHO/BS/2016.2300 – (Page:89 – 34th Line) | |
| Efficiency Analysis | SRID assay /  RT-qPCR | 1. Similar immunodiffusion in fixed dilutions  2. Similar copies | 1-Vaccine raw material viral structures give a positive reaction in dilutions similar to virüs specific antibodies.  2- It must be shown that the number of similar copies is preserved. | Effective | 1.WHO Technical Report Series No. 979, 2013.  2. 20 July 2017  EMA/CHMP/BWP/310834/2012 Rev.1 | |
| Replicative Virus Test | Real-Time Cell Analysis (RTCA) / MTT Based Cytopathic Effect Test | NEGATIVE | Cytopathic effect observation with 3 weeks of VERO cell supernatant. | Negative | (Gorshkov et al., 2020) | |
| Microbiological Quality Control | Culture | NEGATIVE | - | Negative | EP-01/2017:50203 | |
|  | Gram Stain | NEGATIVE | - | Negative | EP-01/2017:50203 | |
| Fungi Culture | Culture | NEGATIVE | - | Negative | 2.6.7 (method reference)  EP-01/2017:50203 | |
| Mycoplasma | PCR | NEGATIVE | - | Negative | 2.6.1 (method reference)  EP-01/2017:50203 | |
| Endotoxin | LAL Test | <5 U/dose | - | Negative | EP-01/2010:50110 (Page:712) | |
| Stability | Single Radial Immunodiffusion (SRID) / RT-qPCR | 1. Similar immunediffusion in fixed dilutions.  2.Less than 0.1% change in RT-qPCR. | Efficiency analysis of 1,2,3,6,12 and 24th months under <-60 ˚C storage conditions. | 1th Month Positive / No Change | 1.WHO Technical Report Series No. 979, 2013.  2. 20 July 2017  EMA/CHMP/BWP/310834/2012 Rev.1 | |
| **COVID-19 VACCINE PRODUCTION END OF PROCESS (BEFORE LYOPHILIZED LAST PRODUCT DISSOLVATION)** | | | | | | |
| **PARAMETERS** | **METHODS** | **LIMITS** | **EXPLANATIONS** | **RESULTS** | **REFERENCES** | |
| Appearance | With Eyes | Dry Grayish Powder | Content, label, and packaging control. | Dry grayish powder.  Suitable content, label, and packaging. | Particle Smudge (Visible Particles)  (2.9.20 Method References)  EP-01/2008:20920 | |
| Relative Humidity | Moisture Analyzer | < %5 | - | The result is expected. | (Biryukov et al., 2020) | |
| **COVID-19 VACCINE PRODUCTION END OF PROCESS (AFTER LYOPHILIZED LAST PRODUCT DISSOLVATION)** | | | | | | |
| **PARAMETERS** | **METHODS** | **LIMIT** | **EXPLANATIONS** | **RESULTS** | **REFERENCES** | |
| Identity | Single Radial Immunodiffusion (SRID) | POZİTİF | Identity with SARS-CoV-2 specific S protein antibody. | The result is expected. | EP-01/2008:1375 | |
|  | RT-qPCR | > 1x10^7^ | Virus copy number / dose analysis. | Active Virus | 1x10^14^ & 1x10^15^ | EP-04/2009:2308 |
|  |  |  |  | Inactivated Virus | 9x10^7^ & 1,2x10^11^ |  |
| Particle Dissolution Rate | With Eyes | ≤ 1 minute | - | 0.5 second | Reconstitution Time < 1 minute  EMA/CHMP/88371/2018 (Page:22 – Table:5) | |
| Biochemical Analysis | pH, Ca, Total Protein, Albumin, Osmolalite | pH:6-8  Ca: <1mg/L  Total protein: <1g/dl  Albumin: <0.6g/dl  Osmolalite: >240 mosmol/kg | - | pH:6,55  Ca: 0,1 mg/dL  Total Protein:  No HSA (10^15^) : 3,45 μg/dose  With HSA (10^15^) : 28,27 μg/dose Albümin: <50ng/dose  Osmolarite: 284 mOsm/kg | Potentiometric determination of pH (2.2.3 method reference):  EP-07/2016:20203  Ca (2.4.3 Calcium):  EP-01/2008:20403  WHO Expert Committee on Biological Standardization  Fifty-sixth report.  WHO Technical Report Series, No. 941 (A.3.3.4 –Page:312 When calcium adjuvants are used, the concentration of calcium should not exceed 1.3 mg per single human dose.)  The albumin content ideally should not exceed 1% of total protein content.  Total Protein-Albumin; WHO/BS/2016.2300 – (Page:89 – 34th Line)  Bovine serum albumin: Maximum 0.65 μg per human dose, determined by an appropriate immunochemical method. (2.7.1).  EP-01/2009:2418  Osmolarity:  EP-01/2012:20235  EP-07/2012:2061 (2.2.35 It should be within the limits approved for each preparation.)  Monoclonal Antibodies for Humans  Osmolality (2.2.35).  Unless justified and permitted, it should be at least 240 mmol / kg.  EP-01/2012:2031 | |
| Purity | VERO Protein ELISA | <50 ng | - | <50 ng | (Jia et al., 2009) | |
|  | VERO DNA NanoDrop | < 10 ng/ml | - | < 10 ng/ml | If a continuous cell line is used for virus replication, the host-cell DNA residue content is determined using an appropriate method, not more than 10 ng at a human dose.  EP-04/2009:2308 | |
|  | Penicillin, Streptomycin, Ciprofloxacin | < %0.01 | - | < %0.01 | Penicillin and streptomycin should not be used at any stage of production or added to the final product; however, master seed batches prepared with a medium containing penicillin or streptomycin can be used for production when proven and validated.  EP-01/2017:0153 | |
| Replicative Virus Test | Real-Time Cell Analysis (RTCA) / MTT Based Cytopathic Effect Test | NEGATIVE | Cytopathic effect observation with 3 weeks of VERO cell supernatant. | Negative | (Gorshkov et al., 2020) | |
| Microbiological Quality Control | Culture | NEGATIVE | - | Negative | 2.6.1 (method references)  Bacterial and fungal contamination.  The master cell bank and each working cell bank comply with the sterility test (2.6.1) using 10 mL of supernatant from cell cultures. The test is done in 1% of the containers, at least 2 containers are used.  EP-01/2017:50203 | |
|  | Gram Stain | NEGATIVE | - | Negative | 2-3-1-5. Biochemical determination based on physiological reactions  Measurement principles: Gram staining or other early discrimination tests are used to decide the appropriate test protocol before these determinations.  EP-07/2017:50106 | |
| Fungi Culture | Culture | NEGATIVE | - | Negative | 2.6.1 (method references)  Bacterial and fungal contamination.  The master cell bank and each working cell bank comply with the sterility test (2.6.1) using 10 mL of supernatant from cell cultures. The test is done in 1% of the containers, at least 2 containers are used.  EP-01/2017:50203 | |
| Mycoplasma | PCR | NEGATIVE | - | Negative | 2.6.7 (method references)  Mycoplasma (2.6.7). The master cell bank and each working cell bank fit the mycoplasma test. One or more containers are used for testing.  EP-01/2017:50203 | |
| Endotoxin | LAL Test | <5 U/dose |  | Negative | 2.6.14 (method references)  Intravenous 5.0 IU endotoxin per kilogram body weight.  P-01/2010:50110 (Page:712-713) | |
| Stability | Single Radial Immunodiffusion (SRID) / RT-qPCR | 1. Similar immunediffusion in fixed dilutions.  2.Less than 0.1% change in RT-qPCR. | Efficiency analysis of 1,2,3,6,12 and 24th months under <-60 ˚C storage conditions. | 1th Month Positive / No Change | 1.WHO Technical Report Series No. 979, 2013.  2. 20 July 2017  EMA/CHMP/BWP/310834/2012 Rev.1 | |
| Activity Analysis | Cytokine Bead Array (CBA) | Cytokine Release >10% relative to negative control | - | > 10% | EP-01 /2008:20724 | |
